# Construction of a 3D whole organism spatial atlas by joint modeling of multiple slices

**DOI:** 10.1101/2023.02.02.526814

**Authors:** Gefei Wang, Jia Zhao, Yan Yan, Yang Wang, Angela Ruohao Wu, Can Yang

**Affiliations:** Department of Mathematics, The Hong Kong University of Science and Technology, Hong Kong SAR, China; Division of Life Science, The Hong Kong University of Science and Technology, Hong Kong SAR, China; Guangdong-Hong Kong-Macao Joint Laboratory for Data-Driven Fluid Mechanics and Engineering Applications, The Hong Kong University of Science and Technology, Hong Kong SAR, China; Department of Chemical and Biological Engineering, The Hong Kong University of Science and Technology, Hong Kong SAR, China; State Key Laboratory of Molecular Neuroscience, The Hong Kong University of Science and Technology, Hong Kong SAR, China; Center for Aging Science, The Hong Kong University of Science and Technology, Hong Kong SAR, China

## Abstract

Spatial transcriptomics (ST) technologies are revolutionizing the way that researchers explore the spatial architecture of tissues. Currently, ST data analysis is often restricted to 2D space within a single tissue slice, limiting our capacity to understand biological processes that take place in 3D space. Here, we present STitch3D, a unified computational framework that integrates multiple 2D tissue slices to reconstruct 3D cellular structures from the tissue level to the whole organism level. By jointly modeling multiple 2D tissue slices and integrating them with cell-type-specific expression profiles derived from single-cell RNA-sequencing data, STitch3D simultaneously identifies 3D spatial regions with coherent gene expression levels and reveals 3D distributions of cell types. STitch3D distinguishes biological variation among slices from batch effects, and effectively borrows shared information across slices to assemble powerful 3D models of tissues. Through comprehensive experiments using diverse datasets, we demonstrate the performance of STitch3D in building comprehensive 3D tissue architectures of the mouse brain, the human heart, and the *Drosophila* embryo, which allow 3D analysis in the entire tissue region or even the whole organism. To gain deeper biological insights, the outputs of STitch3D can be used for downstream tasks, such as inference of spatial trajectories, identification of spatially variable genes enriched in tissue regions or subregions, denoising or imputation of spatial gene expressions, as well as generation of virtual tissue slices.

## Introduction

Spatial transcriptomics (ST) technologies enable high-throughput profiling of gene expressions in intact tissues [1, 2]. ST technologies have been adopted to study organs like hearts and brains as well as the whole organisms like developing embryos from different animal species [3, 4, 5]. Recently, an increasing number of ST datasets consisting multiple parallel 2D slices (*x-y*-axis) along the *z*-axis in a tissue have been generated [6, 7, 8, 9], but most existing ST tools only provide analysis results for one single 2D tissue slice. Biological organs and embryos are organized as complex structures in 3D space, and biological processes rarely occur in a single 2D plane. For example, morphogen gradients spread in 3D space to instruct cell type differentiation and morphogenesis to organize into functional organs during embryogenesis. Having only a 2D view of single tissue slice significantly restricts our interpretation of biological signaling and processes; how they influence organization of different cell types; and how cells interact with each other at the organ level.

With each single slice characterizing a 2D spatial transcriptomic landscape, joint modeling of multiple 2D slices accumulated in recent studies offers opportunities to depict a comprehensive 3D picture of biological systems. Current progresses in jointly handling multiple parallel 2D slices mainly focus on achieving the alignment of spots across tissue slices [10]. Yet comprehensive 3D characterization of tissues or whole organisms requires additional 3D analyses from joint modeling of multiple tissue slices as a whole, so that fine-grained 3D structures in entire tissue regions can be jointly analyzed. This is essential to identify genes with 3D spatial patterns, and to detect 3D cell type gradients in spatial organizations. To this end, additional efforts with joint analysis of multiple tissue slices, to annotate meaningful 3D spatial tissue regions and decipher 3D spatial distributions of cell types in tissues or organs are needed. Specifically, the following two 3D analysis tasks are fundamental. The first task is to identify biologically interpretable 3D spatial regions with spots having similar gene expression patterns and close spatial locations. With this task, meaningful tissue structures which are related to biological functions can be detected. An example of this is the 3D layer organizations of the cerebral cortex. These spatial region annotations then enable meaningful downstream analyses, including spatial trajectory inference among nearby 3D spatial regions and detection of spatially variable genes related to 3D spatial regions. The second task is to infer 3D spatial distributions of fine-grained cell types in tissues by combining ST datasets with abundantly available single-cell RNA-sequencing (scRNA-seq) atlases. Existing ST technologies based on next generation sequencing (seq-based ST approaches), such as 10x Visium, can detect transcriptome-wide gene expression within spatial spots, but each spot often contains multiple cells, resulting in a relatively lower spatial resolution. With utilization of additional information from scRNA-seq atlases, the 3D cell-type deconvolution task decomposes cell-type mixtures in spots and thereby achieves higher resolution for 3D reconstruction of ST data. Such cell-type composition information allows greater insight into the possible biological functions of specific cell-type-enriched regions. Meanwhile, the effective usage of scRNA-seq data not only effectively resolves the 3D spatial organization of fine-grained cell types, but also facilitates the denoising of 3D gene pattern measured by ST data.

Here, we present STitch3D to characterize 3D tissue architectures from multiple tissue slices by addressing the aforementioned two 3D analysis tasks, i.e., identifying 3D spatial regions and revealing 3D cell-type distributions. By joint modeling of gene expression levels and spatial locations from multiple tissue slices, STitch3D is able to distinguish biological variation among slices from batch effects, and effectively integrate information across slices for assembling powerful 3D models of tissues. Moreover, STitch3D simultaneously addresses the 3D spatial domain detection task and the 3D cell-type deconvolution task in a unified framework, providing two connected and complementary views of reconstructed 3D tissue architectures. Compared to existing ST data analysis tools, nearly all of which are developed for analyzing one single tissue slice by performing either the spatial domain detection task [11, 12, 13] or the cell-type deconvolution task [14, 15, 16, 17, 18], STitch3D enables a 3D view of the cellular architecture of tissues with accurate 3D domain detection and 3D cell-type deconvolution results, facilitating various downstream analyses.

Through real data studies, we illustrate STitch3D’s improvements in accuracy gained by borrowing information across multiple slices. Through the application of STitch3D to diverse datasets, we demonstrate STitch3D’s capability to correctly recover spatial regions with biological meanings, and elucidate spatial distributions of fine-grained cell types across ST slices. In particular, STitch3D is also scalable to handle dozens of different tissue slices, for example, it effectively integrated 35 tissue slices for building a 3D cellular structure of the mouse brain. With its effective integration across multiple tissue slices, STitch3D yields reliable 3D reconstruction results at tissue-scale and even for a whole organism. As a demonstration, we reconstructed a 3D *Drosophila* embryo [9] and used the resulting 3D map to annotate subregions of the developing *Drosophila* gut as well as to profile the local gene expression landscape associated with each subregion. The output of STitch3D can also be applied to downstream tasks such as inference of spatial trajectories, identification of spatially variable genes enriched in tissue regions or subregions, denoising or imputation of spatial gene expressions, as well as generation of virtual tissue slices. STitch3D is now a publicly available Python package (https://github.com/YangLabHKUST/STitch3D), serving as an efficient and reliable tool for ST studies.

## Results

### Method overview

STitch3D is a computational method that uses multiple 2D ST tissue slices to reconstruct the comprehensive 3D structure of tissues (Fig. 1). The inputs to STitch3D are spatially resolved gene expression matrices with 2D spatial coordinates from multiple ST slices, and cell-type-specific gene expression profiles from a paired scRNA-seq reference dataset (Fig. 1**a**). By jointly analyzing all the inputs, STitch3D learns to simultaneously detect 3D spatial domains with coherent gene expressions and recover 3D spatial distributions of fine-grained cell types for achieving detailed 3D tissue models (Fig. 1**c**).

**Figure 1:**
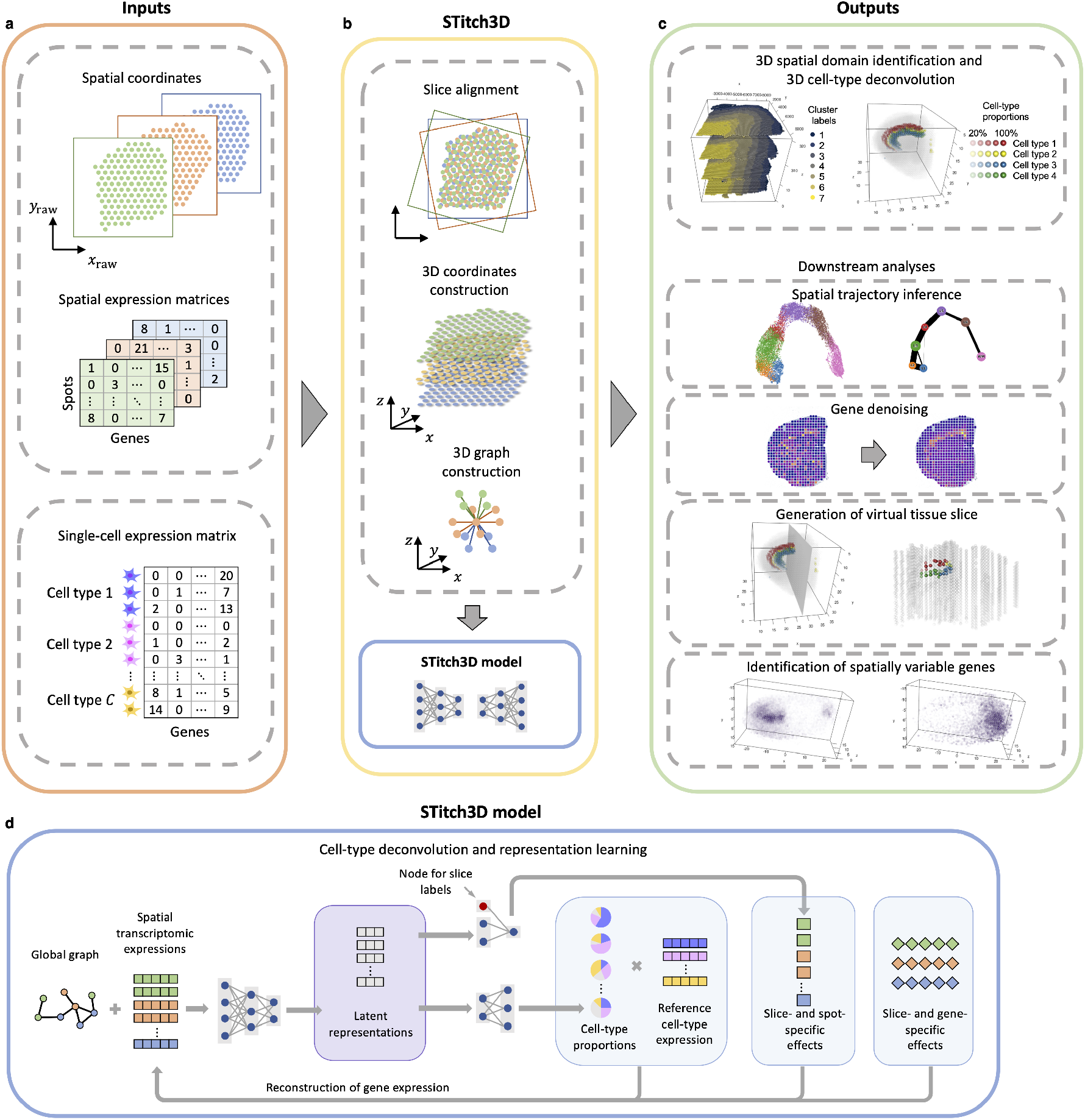
Overview of STitch3D. **a**. Raw data from multiple ST tissue slices, and cell-type-specific gene expression profiles from a reference scRNA-seq dataset are taken as STitch3D’s inputs. **b**. The preprocessing steps of STitch3D include alignment of spots from different tissue slices to build 3D locations of spots, and construction of a global 3D graph. The main model of STitch3D incorporates these structures to perform representation learning for 3D spatial domain identification and 3D cell-type deconvolution. **c**. STitch3D outputs the 3D spatial region identification result and the estimation of 3D spatial distributions of different cell types in tissues. STitch3D also enables multiple downstream analyses, including spatial trajectory inference, denoising of low-quality gene expression measurements, generation of virtual tissue slices, and identification of genes with 3D spatial expression patterns. **d**. STitch3D jointly models multiple slices, and utilizes a graph attention-based neural network to learn latent representations of spots and cell-type proportions with 3D spatial information.

The preprocessing steps of STitch3D include building 3D spatial coordinates for spots by aligning multiple tissue slices, and constructing the global 3D neighborhood graph for spots (Fig. 1**b**). With these inputs, STitch3D then trains a deep learning-based model to effectively integrate information across all tissue slices, and perform both 3D spatial region detection and 3D cell-type decomposition tasks (Fig. 1**d**). For STitch3D, we introduce a shared latent space to distill meaningful biological variations contained in multiple tissue slices and facilitate the removal of batch effects across slices. In the latent space, each spot has its representation which is used to perform the spatial domain identification and cell-type deconvolution tasks jointly.

To be specific, STitch3D uses an encoder network to map spatial spots from multiple slices into the shared latent space, in which spots with similar expression levels and spatial locations tend to have similar representations. We achieve this goal by adopting a graph neural network [19] with the global 3D neighborhood graph for spots in the encoder network. STitch3D then uses another neural network, namely deconvolution network, to infer cell-type proportions from latent representations of spots. The deconvolution network is optimized to reconstruct the raw ST gene expressions by combining the estimated cell-type proportions with the cell-type-specific gene expression profiles from the scRNA-seq reference. With the model innovations, STitch3D can account for potential batch effects across slices and technical effects between ST and scRNA-seq technologies. After training the model, STitch3D learns spatially informed representations of spots and cell-type decompositions of spots. These latent representations are used for 3D spatial domain identification with community detection algorithms. More downstream analyses, e.g., spatial trajectory inference, can also be performed with the spot representations. As cell-type proportions are generated from latent representations of spots, estimated cell-type proportions are also aware of 3D spatial information. Thus, they are able to recover the 3D spatial distribution of cell types in the tissue. Combined with the cell-type-specific gene expression profiles from the scRNA-seq reference, the spatial-aware proportions of cell types can also help to denoise the low-quality spatial measurements of genes, or predict the expression levels of unmeasured genes. Additionally, owing to effective detection of 3D spatial regions and inference of 3D cell-type spatial distributions, STitch3D also enables downstream analyses like identification of spatially variable genes enriched in a specific 3D region and detection of cell type gradients in newly generated virtual slices (Fig. 1**c**). Details are included in the Method section.

### STitch3D’s jointly modeling of multiple slices enables improved spatial domain detection and cell-type deconvolution accuracy

In this section, we first illustrated that although STitch3D is designed for 3D reconstruction of tissues by integrating multiple slices, it can produce accurate spatial domain detection and cell-type deconvolution results when dealing with one single slice. More importantly, we also showed that STitch3D’s multi-slice analysis, which jointly models multiple slices, leads to further improved performance in both spatial domain detection and cell-type deconvolution, compared to that of STitch3D’s single-slice analysis.

We first compared STitch3D’s single-slice and multiple-slice results in terms of spatial domain detection performance. For the comparison, we used a human dorsolateral prefrontal cortex (DLPFC) dataset run on the 10x Visium platform [7]. The dataset contains four DLPFC tissue slices with slice indices 151673-151676, collected from the same neurotypical adult donor. Six DLPFC layers (L1-L6) and white matter (WM) were manually annotated for each single slice by researchers based on cytoarchitecture and gene markers [7]. We first applied STitch3D on each single slice to show its effectiveness. Based on the results, we found that in single-slice experiments, STitch3D stably recovered layer structures for all the four slices with its spatial domain detection function (Supplementary Fig. 1). When applied to multi-slice analysis, STitch3D yielded more consistent results across the four analyzed tissue slices, facilitating the 3D reconstruction of the tissue (Fig. 2**a**). Specifically, combining the aligned 3D spatial locations with the consistently detected domains, STitch3D enabled identification of DLPFC layer structures in 3D view (Fig. 2**b**). STitch3D’s 3D spatial domain detection result has very similar pattern compared to manual annotations, indicating its high reliability (Fig. 2**b**, **c**). To quantitatively evaluate the performance, we considered the manual annotations as ground truth, and assessed the accuracy of compared methods based on adjusted Rand index (ARI). A higher ARI score indicates higher similarity between the result and the ground truth. As shown by relatively higher ARI scores, better accuracy was achieved in STitch3D’s multi-slice result compared to its single-slice result, indicating that STitch3D can effectively borrow information across slices for identifying 3D spatial domains (Fig. 2**d**). Meanwhile, the integrated representations of spots in STitch3D’s shared latent space also show clear spatial trajectory from L1 to L6 and WM (Fig. 2**e**).

**Figure 2:**
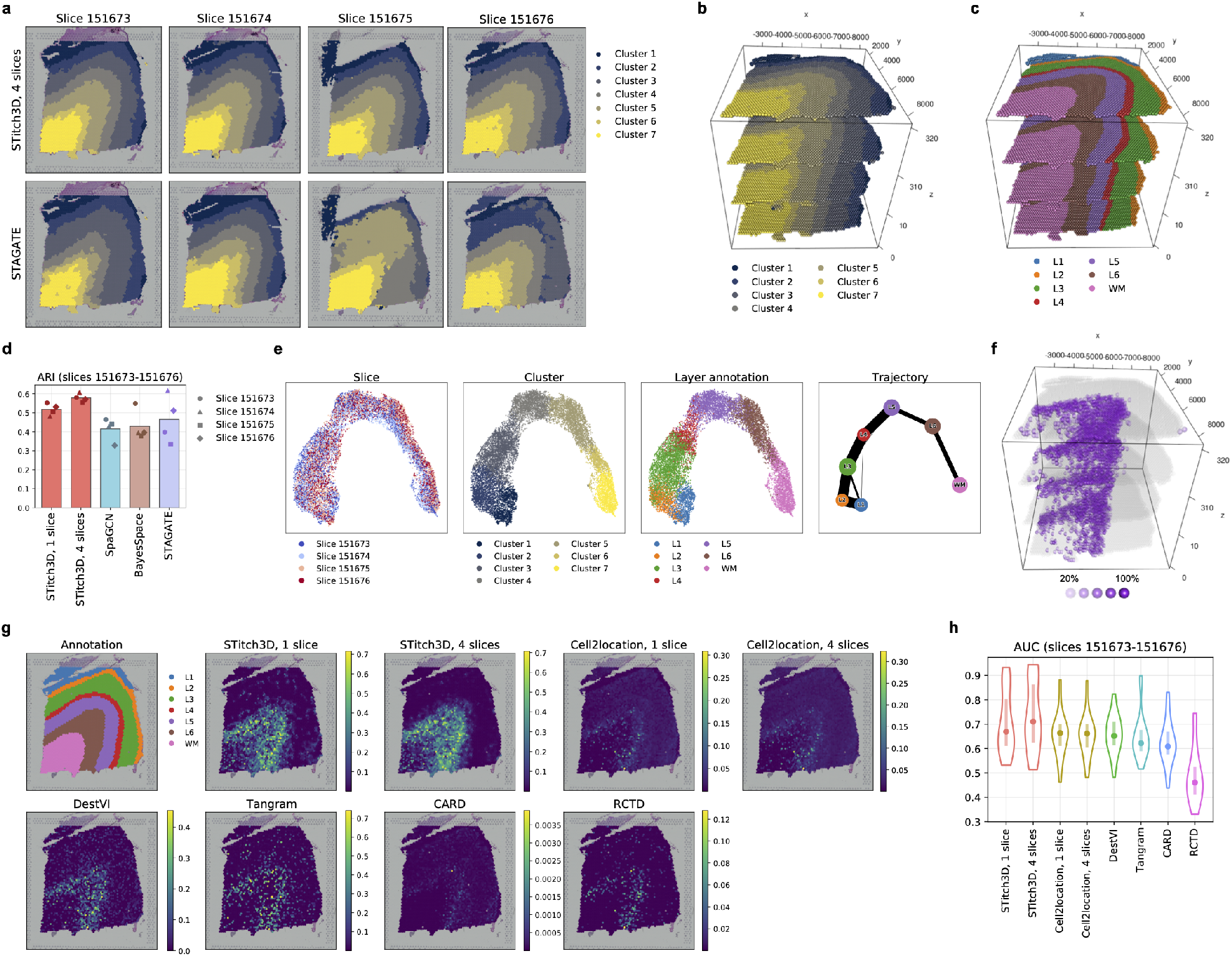
3D spatial domain detection and cell-type deconvolution for the DLPFC dataset. **a**. STitch3D’s and STAGATE’s spatial domain detection results on the four tissue slices. **b**. STitch3D’s 3D spatial domain detection result. **c**. 3D visualization of manual annotations. **d**. ARI scores of spatial domain detection methods, which measure the similarity between assigned spatial domain labels and manual annotations. **e**. We visualized STitch3D’s learned representations of spots in the shared latent space with UMAP plots [20], colored by slice indices, STitch3D’s assigned domain labels and manual annotations (three panels on the left). With the representations of spots, the PAGA algorithm [21] was adopted for spatial trajectory inference (the rightmost panel). f. STitch3D’s reconstructed 3D spatial distribution of Ex_8_L5_6 neuronal subtype. Spots with proportion values larger than 20% are shown in purple color, and a lower transparency indicates a higher proportion. **g**. Estimated Ex_8_L5_6 neuronal subtype proportions from cell-type deconvolution methods on slice 151676. **h**. Violin plots of AUC scores of compared methods measured with ten neuronal subtypes.

Next, we compared the performance of STitch3D’s single-slice and multiple-slice results in terms of cell-type deconvolution. Throughout the comparison conducted with the DLPFC Visium data, we used a single-nucleus RNA-sequencing (snRNA-seq) dataset of human DLPFC [22] as the cell-type gene expression reference. In the reference dataset, ten excitatory neuronal subtypes were identified. The layer specificity of each neuronal subtype was determined based on the subtype’s expression patterns of the layer markers. We assessed the reliability of STitch3D’s cell-type deconvolution results based on the receiver operating characteristic (ROC) analysis, in which the estimated proportions of neuronal subtypes were used to recover the layer specificity. A higher value of area under curve (AUC) of ROC indicates more reliable performance. As shown in Fig. 2**h**, the accuracy scores obtained in STitch3D’s multi-slice analysis became higher than the accuracy scores of STitch3D’s single-slice results. Besides quantitative evaluation, we also visualized cell-type deconvolution results (Supplementary Fig. 2). Take the neuronal subtype named Ex_8_L5_6 as an example. This subtype has high expression levels of marker genes of layers L5 and L6. STitch3D showed the clear enrichment of this neuronal subtype in layers L5 and L6 (Fig. 2**g**). Consistent with quantitative evaluation, we observed decreased and less noisy Ex_8_L5_6 proportions in layers L1-L4 in STitch3D’s multi-slice result, compared to single-slice analysis. With multi-slice analysis, STitch3D’s 3D cell-type deconvolution result on Ex_8_L5_6 neuronal subtype and other neuronal subtypes clearly characterized their 3D spatial distributions (Fig. 2**f**).

To show the advantage of STitch3D, we compared STitch3D with several existing representative ST data analysis tools, most of which were developed to deal with only single slice. For the spatial domain detection task, we evaluated methods including BayesSpace [11], SpaGCN [12] and STAGATE [13]. However, unlike STitch3D’s single-slice and multi-slice results, these methods showed limited capability of detecting cortical layers for at least one tissue slice (Fig. 2**a** and Supplementary Fig. 1). Their relatively less satisfactory performances were also indicated by lower ARI scores (Fig. 2**d**). We then compared STitch3D with cell-type deconvolution methods including RCTD [14], Cell2location [15], Tangram [16], DestVI [17] and CARD [18]. In single-slice experiments, STitch3D showed higher AUC scores compared to all the other methods (Fig. 2**h**). Among these methods, Cell2location also allows for multi-slice analysis, so we applied it to analyze four DLPFC tissue slices jointly. Nevertheless, being different from STitch3D, Cell2location showed limited ability to borrow information across slices, as indicated by its similar AUC scores in single-slice and multi-slice experiments (Fig. 2**h**). For a more comprehensive comparison, we also followed a recent study [23] to assess STitch3D and other cell-type deconvolution methods in three different scenarios. We used datasets profiled by seqFISH+ [24], STARmap [25] and MERFISH [26] with corresponding cell type expression references [27, 26]. As shown by the results in Supplementary Figs. 3, 4 and 5, in single-slice setting, STitch3D showed the best overall performance among all methods. In multi-slice analyses under different scenarios, STitch3D achieved even higher accuracy by leveraging information from adjacent slices, indicating its advantage in joint modeling of multiple slices. More details can be found in Supplementary Sections 1 and 2.

### STitch3D reconstructs the complex 3D spatial organization of the adult mouse brain and allows for analysis of virtual slices

As an organ that controls animal behaviors, the brain has very complex structures and contains diverse and specialized cell types, especially neurons [28]. Therefore, in brain research, it is essentially important to explore the complex anatomical and functional architecture of the brain, and to decipher the detailed organization of diverse neuron types. Such exploration could help researchers characterize structure-function relationship contained in brain and have a better understanding of brain mechanisms.

In this section, we demonstrated that STitch3D is able to accurately reconstruct complex 3D structure of the adult mouse brain by determining brain regions with similar gene expressions and depicting structural architectures of different cell types. Based on the constructed 3D organization of the mouse brain, STitch3D further enables to generate virtual slices, presenting new views of the brain that are not captured by the original slices. In this task, we used 35 coronal slices from one mouse brain hemisphere across the antero-posterior (AP) axis collected from two adult mice [6]. To achieve a fine-grained characterization of brain structure, we adopted a mouse brain snRNA-seq dataset that contains 59 distinct cell types and subtypes [15] as the cell-type expression reference. The reconstruction of the mouse brain with 3D spatial region detection and 3D cell-type deconvolution can be very challenging, because it requires methods to not only account for batch effects contained in dozens of slices, but also to be capable of distinguishing subtle difference among nuanced cell types and subtypes.

When applied to this task, STitch3D successfully reconstructed the 3D mouse brain model by reliably identifying 3D spatial regions and inferring 3D cell-type distributions. First, STitch3D partitioned the mouse brain into 11 well-organized and biologically meaningful 3D spatial regions, based on its integrated latent representations of spots in the shared latent space (Supplementary Fig. 6). For instance, three layer-structured spatial domains detected by STitch3D with cluster labels 1, 2 and 5 together formed the isocortex region (Fig. 3**a**-**c**). Of note, although these slices are all coronal mouse brain sections, they carry important biological difference, because the brain structure gradually changes along the AP axis. In STitch3D’s learned representations of spots in the shared latent space, such variation was well-preserved, instead of being corrected as batch effects (Supplementary Fig. 6). Therefore, STitch3D can characterize regions that vary along the AP axis. For example, cluster 3 in its detected spatial domains corresponded to the hippocampus region (Supplementary Fig. 6), and cluster 9 corresponded to the thalamus region (Supplementary Figs. 6 and 7). Such results demonstrated STitch3D’s ability to identify spatially coherent tissue regions from dozens of ST slices that were not created as “technical replicates” but contain certain biological variation.

**Figure 3:**
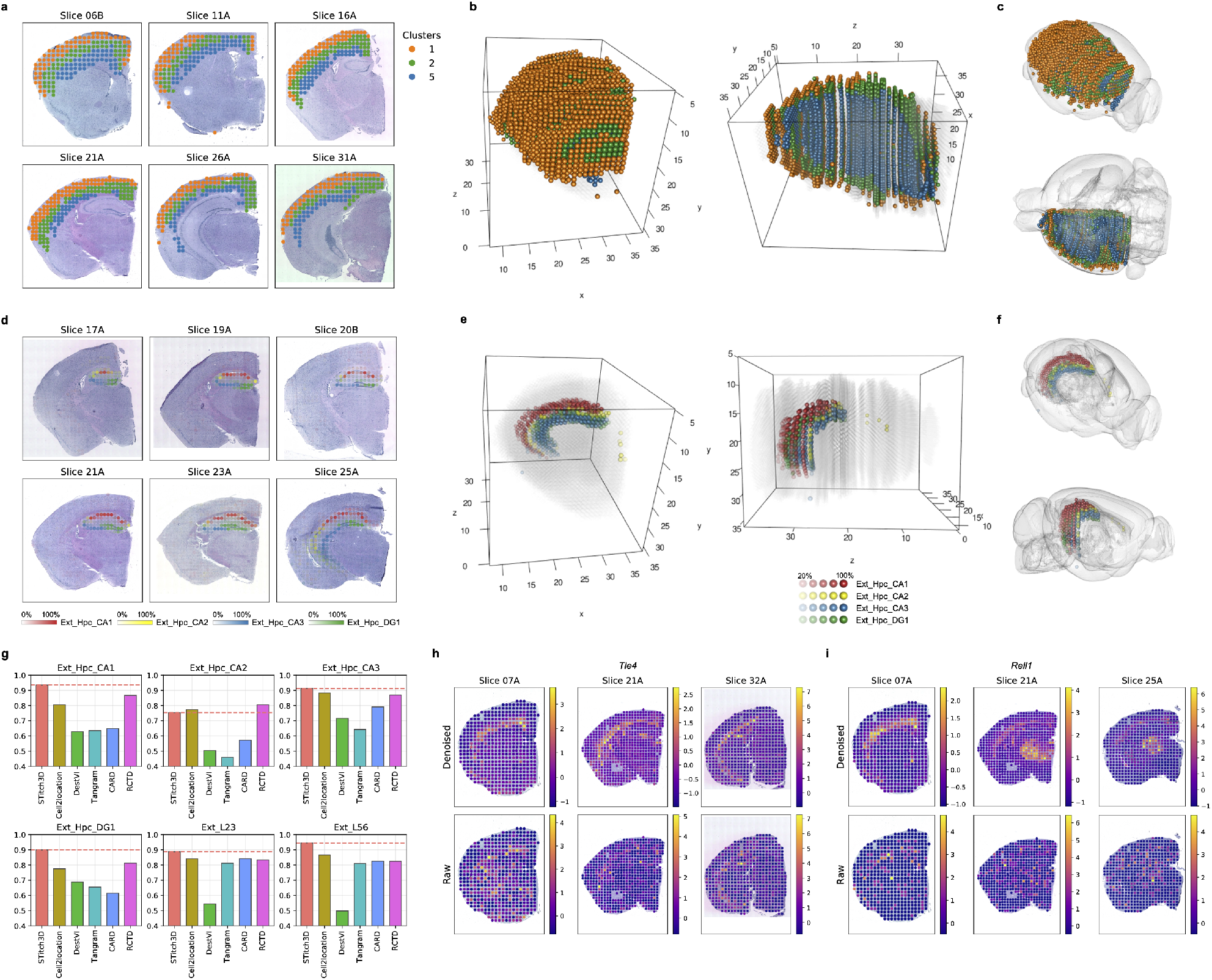
3D reconstruction of the adult mouse brain. STitch3D reconstructed the 3D model of the adult mouse brain with 35 coronal slices. **a**. Clusters 1, 2 and 5 in STitch3D’s spatial domain detection result visualized on 2D ST slices. **b**, **c**. 3D visualizations of clusters 1, 2 and 5 in STitch3D’s aligned 3D coordinates using ICP (**b**) and in the Allen Mouse Brain Common Coordinate Framework (CCFv3) [29] (**c**). **d**. Estimated proportions of four hippocampal neuron types in STitch3D’s cell-type deconvolution result visualized on 2D ST slices. A lower transparency indicates a higher proportion. **e**, **f.** 3D visualizations of proportions of four hippocampal neuron types in STitch3D’s aligned 3D coordinates using ICP (**e**) and in CCFv3 (**f**), where spots with proportion values larger than 20% are shown. **g**. AUC scores of compared methods measured with six regional cell types. **h**, **i**. Denoised and raw expression levels of genes *Tle4* (**h**) and *Rell1* (**i**) in z-scores.

Leveraging cell-type signatures from the snRNA-seq reference dataset containing fine-grained annotations, STitch3D also revealed 3D distributions of specific cell types. As an example, it accurately reconstructed curve-shaped distributions of four hippocampal neuron types in three cornu ammonis (CA) areas and dentate gyrus (DG) (Fig. 3**e**, **f**). Comparing cell-type proportions of these cell types on 2D slices (Fig. 3**d**) with 2D coronal reference provided by the Allen Reference Atlas – Mouse Brain [30] (Supplementary Fig. 8), we found that STitch3D’s reconstructed distributions of these hippocampal neuron types correctly matched corresponding hippocampus areas including CA1, CA2, CA3 and DG. Besides, STitch3D also captured distributions of excitatory neurons across cortex layers, such as excitatory neurons enriched in layers 2-3 and layers 5-6 (Supplementary Fig. 9), and other major regional subtypes (Supplementary Fig. 10). To validate the correctness of STitch3D’s cell-type deconvolution, we used estimated proportions of four hippocampal neuron types to recover CA1, CA2, CA3 and DG regions and compared the result with region annotations in these slices using ROC analysis. Similar ROC analysis was also conducted with the two cortical neuron types in layers 2-3 and layers 5-6. In ROC analyses, we also compared STitch3D with other cell-type deconvolution methods, where STitch3D and Cell2location were applied in multi-slice setting, and the others were tested in single-slice setting. As indicated by the highest AUC values of ROC analyses, STitch3D achieved the best consistency between estimated cell-type proportions and region annotations for five out of six cell types (Fig. 3**g**). It also showed comparable performance with Cell2location and RCTD for the remaining one cell type. The ROC analyses demonstrated that, STitch3D is able to reliably recover distributions of cell types in expected regions.

One remarkable problem in ST studies is that gene expressions measured by ST technologies often suffer from extensive noise, which brings difficulties for studying spatial expression patterns of genes [31]. Thus, it is often necessary to remove such noise of ST data. Combining estimated cell-type proportions with cell-type expression signatures from the reference dataset, STitch3D also enables denoising expression levels of low-quality genes. We demonstrated this application of STitch3D with two genes that had noisy spatial expression patterns in the original dataset, including *Tle4* and *Rell1* (Fig. 3**h**, **i**). After applying STitch3D, the distribution patterns of these two genes became more clear (Fig. 3**h**, **i**). Comparing with references of these genes from *in situ* hybridization (ISH) images and signal intensity heat maps provided by the Allen Mouse Brain Atlas [32] (Supplementary Fig. 11), we found that STitch3D’s denoised genes were highly consistent with spatial expression patterns in the references, demonstrating STitch3D’s reliability in its application to the gene denoising task.

As STitch3D reconstructs the 3D architecture of a tissue, it is able to further provide users with auxiliary views of tissues which are not captured by original slices. This functionality of STitch3D helps us have a deeper understanding of a tissue from different angles. In this example, with the 3D mouse brain model reconstructed by STitch3D using coronal slices, we created a virtual sagittal slice of the mouse brain, by introducing a plane parallel to the AP axis and projecting adjacent spots onto the plane (Fig. 4**a**, **c**). With its estimated cell-type proportions, such as excitatory neurons in hippocampus and thalamus, STitch3D also revealed the mouse brain structure on the sagittal slice with the distributions of different cell types (Fig. 4**b**, **d**). This result is consistent with the sagittal slice reference provided by the Allen Reference Atlas [30] (Supplementary Fig. 12), indicating STitch3D’s correctness in the 3D reconstruction and virtual slice generation.

**Figure 4:**
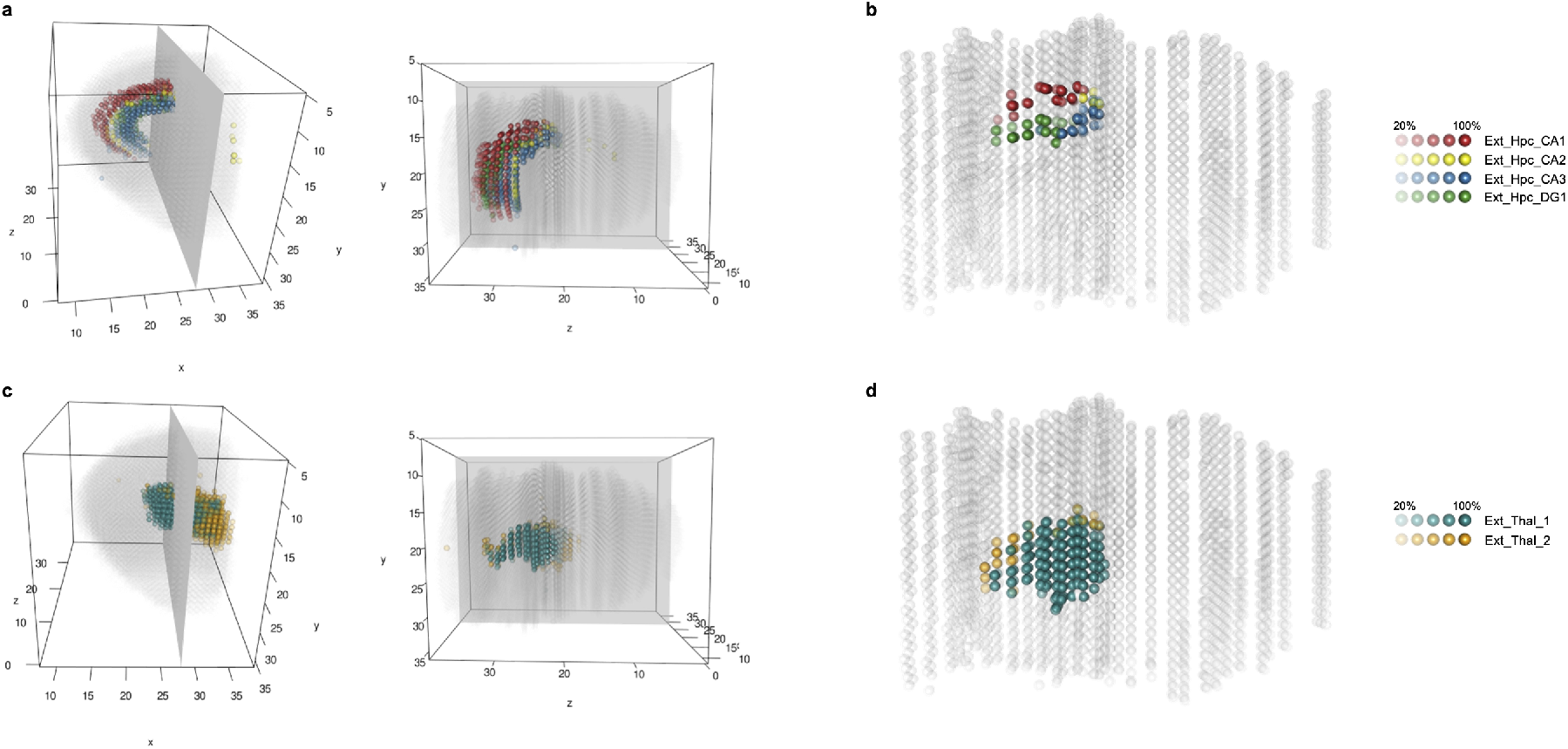
A virtual sagittal slice of the mouse brain created by STitch3D from its 3D reconstruction result. **a**. 3D visualizations of proportions of four hippocampal neuron types and the plane containing the virtual sagittal slice. **b**. The virtual sagittal slice with proportions of four hippocampal neuron types. **c**. 3D visualizations of proportions of two thalamic neuron types and the plane containing the virtual sagittal slice. **b**. The virtual sagittal slice with proportions of two thalamic neuron types. In **a**-**d**, spots with proportion values larger than 20% are shown, and a lower transparency indicates a higher proportion.

### STitch3D reconstructs the developing human heart and facilitates spatiotemporal analysis

The heart is the first functional organ to form in the human embryo [34]. Development of the human heart is a complex process that is incompletely understood. Gaining deeper insights into this process requires comprehensive 3D characterizations of gene expression landscape and detailed 3D maps of cell types in the human embryonic hearts. In this section, we showed that STitch3D is capable of building 3D maps of the developing human heart at different time points, facilitating spatiotemporal analysis of the developing human heart.

Recently, a study generated a spatiotemporal dataset describing the gene expression pattern during human heart development using ST [8]. Specifically, multiple tissue slices were collected from the human hearts along the dorsal-ventral axis at three developmental stages, including 4.5-5, 6.5, and 9 post-conception weeks (PCWs), respectively. We used STitch3D to analyze the ST data with the scRNA-seq data obtained from the same study. We first focused on the application of STitch3D on the 6.5 PCW human heart. STitch3D identified five clusters in its shared latent space where nine 6.5 PCW tissue slices were integrated. We found that the five clusters are consistent across multiple tissue slices, though the tissue slices vary along the dorsal-ventral axis, showing STitch3D’s effectiveness in across-slice modeling (Fig. 5**a**). As shown in Fig. 5**b**, the five clusters correspond to anatomical regions in the human heart. For example, the outflow tract (OFT) was detected by cluster 3. The left atrium and the right atrium were identified as cluster 4. With the scRNA-seq atlas providing cell-type-specific gene expression profiles, STitch3D also generated 3D spatial distributions of fine-grained cell types in the human embryonic heart, which is essential in establishing a 3D atlas. First, it correctly identified distributions of three different cardiomyocyte subtypes. In its cell-type deconvolution result, atrial cardiomyocytes and ventricular cardiomyocytes were found to be specifically localized in the atria (cluster 2) and ventricles (clusters 1, 2 and 5), respectively (Fig. 5**c**, **d**, **j**). Meanwhile, *Myoz2*-enriched cardiomyocytes were found in both the atria and ventricles (Fig. 5**c**, **j**). Such result was confirmed by the original study [8], in which a human heart section was partitioned into the atrium region and the ventricle region, and cell-type compositions in these two parts were examined by scRNA-seq respectively. Besides cardiomyocytes, we found that smooth muscle cells have a considerable rich density in the OFT region (cluster 3) (Fig. 5**c**, **e**, **j**), which is consistent with the expression pattern of smooth muscle-related marker gene *MYH11* (Supplementary Fig. 13). Based on STitch3D’s result, we also observed that the 6.5 PCW human heart is surrounded by epicardium-derived cells (EPDCs) (Fig. 5**e**). This result is supported by the expression pattern of EPDC marker gene *TBX18* (Supplementary Fig. 13). STitch3D’s cell-type deconvolution result also revealed spatial colocalization relationship between cell types. As indicated by the correlation of cell-type distributions (Fig. 5**h**), atrial cardiomyocytes and ventricular cardiomyocytes distribute in separate areas, but they are all colocalized with *Myoz2*-enriched cardiomyocytes. Besides, as two endothelial cell types, one cell type annotated as capillary endothelium is colocalized with ventricular cardiomyocytes, while the other annotated as endothelium / pericytes is mainly colocalized with EPDCs, indicating the difference in locations of these two similar cell types.

**Figure 5:**
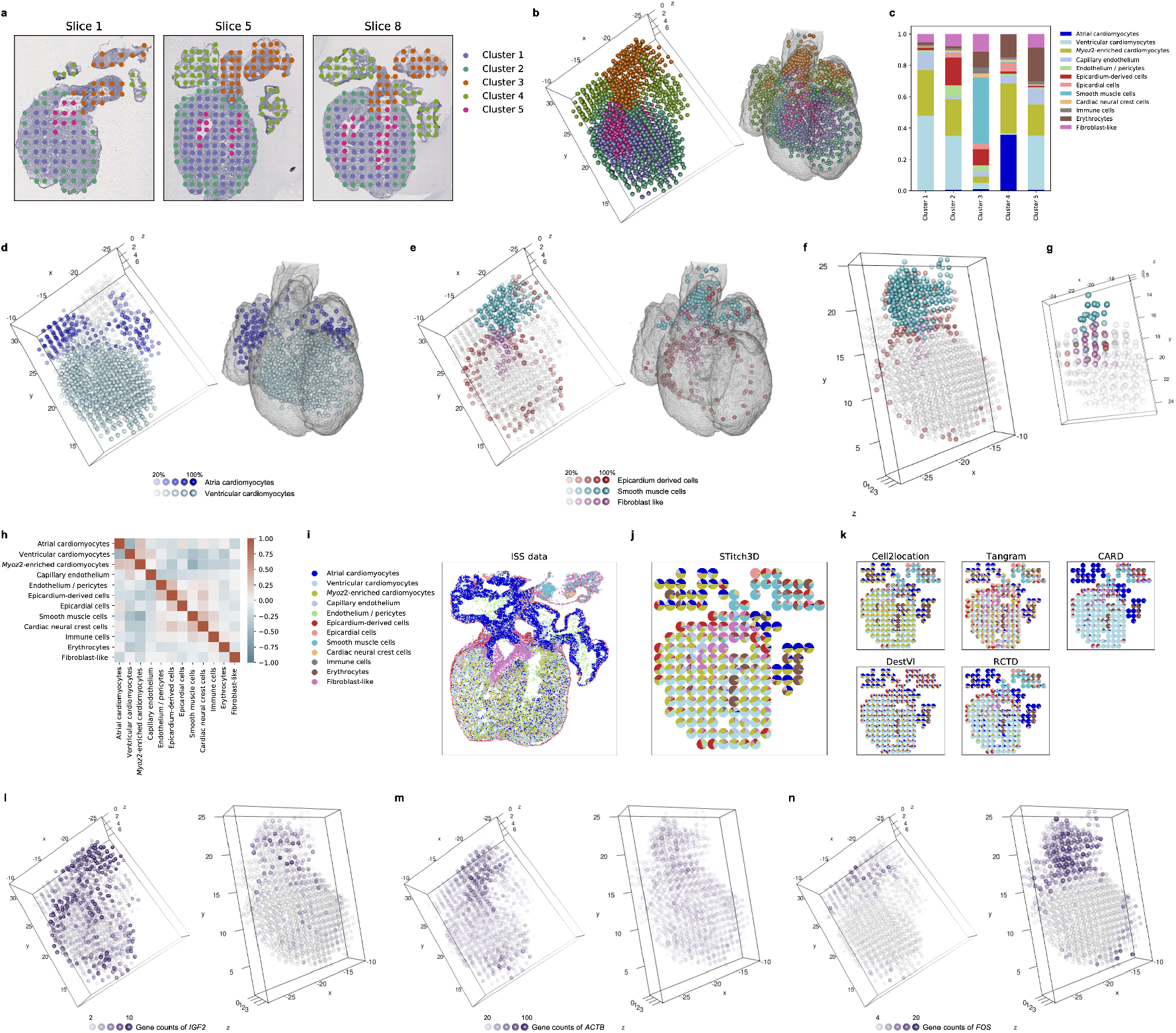
3D reconstruction of the developing human heart. We applied STitch3D to construct 3D atlases of the 4.5-5, 6.5, and 9 PCW human embryonic hearts. **a**. STitch3D’s spatial domain detection result visualized on 2D ST slices from the 6.5 PCW heart. **b**. The detected domains visualized in STitch3D’s aligned 3D coordinates of the 6.5 PCW slices, and in a heart model of Carnegie stage 18 embryo (CS18-6524) [33], respectively. **c**. Based on STitch3D’s result, we visualized the average proportions of cell types in the spatial domains. **d**. 3D spatial distributions of atrial cardiomyocytes and ventricular cardiomyocytes in STitch3D’s aligned 3D coordinates of the 6.5 PCW slices, and in CS18-6524, respectively. **e**, **f**, **g**. 3D spatial distributions of EPDCs, smooth muscle cells, and fibroblast-like cells in the 6.5 PCW heart (**e**), the 9 PCW heart (**f**) and the 4.5-5 PCW heart (**g**). **h**. Spatial colocalization of cell types in the 3D atlas of the 6.5 PCW heart constructed by STitch3D. **i**. The ISS data of a 6.5-7 PCW heart slice with cell-type map created by the original study. **j**, **k**. Estimated proportions of cell types obtained from STitch3D (**j**) and other compared methods (**k**) in a 6.5 PCW heart slice visualized by pie charts. **l**, **m**, **n**. Spatial expression patterns of genes *IGF2* (**l**), *ACTB* (**m**), and *FOS* (**n**).

We further verified STitch3D’s cell-type deconvolution result with a tissue slice obtained using *in situ* sequencing (ISS) provided in the original study [8] (Fig. 5**i**). We compared cell-type deconvolution methods with the ISS data using the seventh slice from the 6.5 PCW slices (Fig. 5**j**, **k**). Compared to STitch3D, other cell-type deconvolution methods showed less satisfactory performance. For example, Tangram found cardiac neural crest cells distributed all over the tissue slice, but they should only present in the OFT [8]. Cell2location, CARD, DestVI and RCTD underestimated the enrichment of fibroblast-like cells in the OFT. Additionally, CARD did not capture *Myoz2*-enriched cardiomyocytes in this slice, and DestVI did not identify smooth muscle cells in the OFT.

After verifying the reliability of STitch3D’s 3D atlas of the 6.5 PCW human embryonic heart, we then applied STitch3D on the remaining two sets of slices from the 4.5-5 PCW and 9 PCW hearts respectively. Based on STitch3D’s results, we observed an overall conserved pattern of cell-type spatial distributions within the studied time window (4.5-9 PCW) (Fig. 5**j** and Supplementary Fig. 14). For instance, across the three different developmental stages of human heart, the relative locations where atrial cardiomyocytes, ventricular cardiomyocytes, smooth muscle cells and fibroblast-like cells have rich densities are consistent. The result suggested that, before 4.5-5 PCW, the human embryonic heart has established a global spatial pattern of gene expressions and cell-type distributions. This global pattern is maintained in later developmental process of heart. In contrast, detailed temporal differences during human cardiac development were also observed, based on the comparison among the 3D atlases across time stages. EPDCs were found to be enriched on the surfaces of the 6.5 PCW and the 9 PCW embryonic hearts, while a considerable lower density of EPDCs was observed on the surface of the 4.5-5 PCW heart (Fig. 5**f**, **g** and Supplementary Figs. 13, 15 and 16). Meanwhile, compared to late stages, the 4.5-5 PCW heart presented a higher density of epicardial cells in its surrounding layer (Fig. 5**j** and Supplementary Fig. 14). Next, we conducted further analysis based on STitch3D’s results to investigate differences between the 6.5 PCW and the 9 PCW embryonic hearts. As we had observed that EPDCs were detected on surfaces of both the 6.5 PCW and the 9 PCW embryonic hearts, here we focused on examining changes of gene expression levels in the EPDC-enriched spots between the two embryonic hearts. To this end, we performed t-test on gene expression levels in EPDC-enriched spots detected by STitch3D, and found that EPDC-enriched spots in the 6.5 PCW embryonic heart showed higher expression levels of gene *IGF2* (Fig. 5**l**). We performed similar comparative analyses in smooth muscle cell-enriched spots identified by STitch3D, which mainly concentrated in the OFT region, and found genes *ACTB* and *FOS* expressed differently at these two stages (Fig. 5**m**, **n**). Particularly, *ACTB* showed higher expression levels in smooth muscle cell-enriched spots in the 6.5 PCW embryonic heart, while *FOS* was found to have higher expression levels in the 9 PCW embryonic heart. These nuanced temporal differences identified using STitch3D help to better characterize the process of human cardiogenesis.

### STitch3D establishes a comprehensive 3D whole organism atlas of the *Drosophila* embryo

*Drosophila melanogaster* has been a prominent model organism for deciphering complex developmental mechanisms for over a century. *Drosophila* embryo scRNA-seq have recently become available as resources [35]. New analytical tools are necessary to release the full potential of these datasets and yield new mechanistic insights in embryogenesis processes.

In this section, we applied STitch3D to reconstruct a 3D *Drosophila* embryo model using multiple ST slices profiled by Stereo-seq from a recent study [9]. Specifically, we analyzed the 16-18 h *Drosophila* embryo dataset, in which spatial bins were constructed by merging 20×20 DNA nanoballs into each 10 μm×10 μm-sized bin in the original study. To apply STitch3D in this task, we built a cell-type reference in a scRNA-seq *Drosophila* embryo atlas [35] at the same developmental stage. The cell type annotation is based on expression patterns of known marker genes provided by the Berkeley *Drosophila* Genome Project (BDGP) [36, 37] and FlyBase [38] (Supplementary Table 1 and Supplementary Figs. 17 and 18).

Compared to other tasks in which STitch3D was examined, 3D reconstruction of the *Drosophila* embryo is confronted with an additional challenge: the *Drosophila* embryo ST dataset is highly sparse, as more than 95% genes in most spatial bins have zero count (Supplementary Fig. 19). However, despite the difficulty, STitch3D successfully reconstructed *Drosophila* embryo with high fidelity by integrating the ST dataset with the cell-type reference. For example, in our STitch3D reconstruction, central nervous system (CNS) spots assembled into a spatial structure that recapitulates the embryonic CNS morphology and position in the embryo [39] (Fig. 6**a**). The reconstruction result is consistent with CNS-specific marker gene such as *Obp44a* (Fig. 6**b**). Another example is the successful reconstruction of salivary gland, which again fully recapitulates the morphology and position of salivary gland in the embryo (Fig. 6**d**), and showed similar spatial pattern compared to salivary gland-specific gene marker such as *CG14453* (Fig. 6**e**). Notably, in comparison with all other cell-type deconvolution methods, STitch3D’s result achieved the highest correlation between regional cell-type distributions and marker genes related to the regions (Fig. 6**c**, **f**).

**Figure 6:**
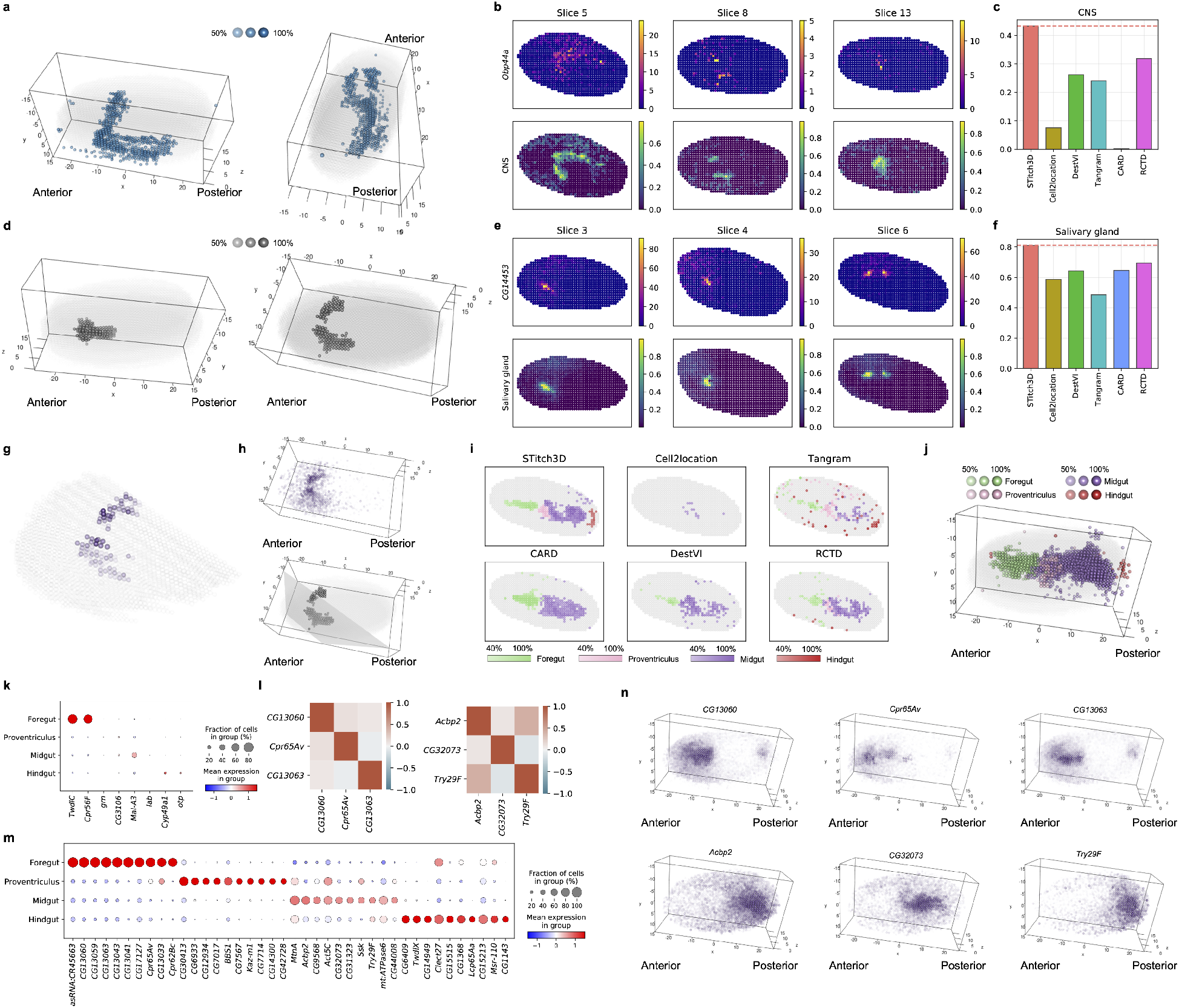
STitch3D constructed the 3D atlas of the *Drosophila* embryo with the Stereo-seq data. **a**, **d**. Visualizations of STitch3D’s estimated 3D distributions of CNS (**a**) and salivary gland (**d**). **b**, **e**. Visualizations of expression pattern of CNS-related marker gene *Obp44a* (**b**) and salivary gland-related marker gene *CG14453* (**e**). We compared the expression patterns of marker genes with STitch3D’s estimated proportions of CNS (**b**) and salivary gland (**e**). **c**, **f.** For all compared methods, we measured Pearson correlations between estimated proportion of cell types (CNS, salivary gland) and pattern of markers (*Obp44a, CG14453*). **g**. We visualized expression levels of *CG14265* in salivary gland-enriched spots on the virtual slice generated by STitch3D. **h**. 3D visualization of spatial expression pattern of *CG14265* (top), and 3D visualization of salivary gland-enriched spots and the plane containing the virtual slice (bottom). **i**. We examined the capability of compared methods to identify foregut, proventriculus, midgut, and hindgut of the *Drosophila* embryo. **j**. Visualization of STitch3D’s estimated 3D distributions of foregut, proventriculus, midgut, and hindgut. **k**. Sparse expression pattern of the known markers in the spatial dataset. **l**. Spatial correlations between STitch3D’s identified markers of foregut and midgut. **m**. We identified marker genes based on STitch3D’s result on the spatial dataset. **n**. Subregional expression pattern of STitch3D’s identified genes of foregut and midgut.

The STitch3D reconstructed embryo then allows us to zoom in to specific regions within an organ type and uncover novel gene expression pattern from new angles in virtual slices. For example, in the salivary gland, we identified that gene *CG14265* exhibited a medial-to-lateral expression gradient (Fig. 6**h**), which was not captured by any of the original slices. Another example is the embryo gut. In STitch3D’s result, foregut, midgut and hindgut were detected in clear coherent spatial regions from anterior to posterior, as well as proventriculus at the junction of the foregut and midgut (Fig. 6**i**, **j** and Supplementary Fig. 20). Notably, other existing methods such as Cell2location, CARD, DestVI and RCTD failed to detect proventriculus or hindgut using the same dataset. Although Tangram detected these four regions in the *Drosophila* embryonic gut, these spots failed to spatially assemble into a coherent structure (Fig. 6**i**). We first visualized the expression levels of known marker genes which were used to annotate the four regions in the scRNA-seq reference. Based on Fig. 6**k**, known foregut-related marker genes *TwdlC* and *Cpr56F*, which highly expressed in the foregut cell population in the scRNA-seq dataset (Supplementary Fig. 18), also showed high expression levels in STitch3D’s detected foregut region in the ST dataset, again validating the accuracy of our reconstruction. Meanwhile, due to the sparsity of this ST dataset, the measured expression levels of known markers of proventriculus, midgut and hindgut in the ST dataset was much lower compared to the expression levels in the scRNA-seq dataset (Fig. 6**k** and Supplementary Figs. 18 and 19). We then used our method to identify novel gut region-specific markers (Fig. 6**m**). Moreover, we are able to further identify subregions within foregut, midgut and hindgut. For example, we detected that *CG13060* is broadly expressed in the foregut, while genes *Cpr65Av* and *CG13063* are highly expressed in subregions of the foregut, indicated by their spatial correlations and 3D distributions (Fig. 6**l**, **n** and Supplementary Fig. 21). Similar findings were observed for the midgut. For example *Acbp2* expresses broadly in the midgut, while *CG32073* and *Try29F* only express in restricted regions in the midgut (Fig. 6**l**, **n** and Supplementary Fig. 22). Notably, the identification of gene patterns in different gut sub-regions will allow biologists to build a comprehensive gene regulatory network that controls the gut differentiation process. To summarize, the above results highlight the power of STitch3D in reconstructing a virtual embryo with accurate regional cell-type distribution patterns in 3D space, which will provide a powerful tool to study a wide range of embryogenesis processes at a systemic level.

## Discussion

In this paper, we have presented STitch3D, an efficient and versatile tool powered by advanced deep learning technologies for 3D reconstruction of tissue structures. By integrating multiple parallel 2D tissue slices, STitch3D is able to characterize 3D architectures, with scales ranging from tissues to whole organisms. With STitch3D’s effective reconstruction of 3D tissue structures through 3D spatial domain detection and 3D cell-type deconvolution, we can identify 3D tissue regions and 3D cell-type distributions without relying solely on manual annotation based on H&E-stained histological images. Using STitch3D’s outputs, we are able to explore the entire 3D landscape of tissue regions and related biological processes in the 3D space. STitch3D’s outputs further enable various downstream analyses, such as identifying genes enriched in a specific 3D region and visualizing cell type gradients in virtual slices that are on different planar sections to the real tissue slices. We have extensively tested STitch3D on diverse datasets including data from human dorsolateral prefrontal cortex, mouse brain, human heart and *Drosophila* embryo, to demonstrate its effectiveness and reliability. On the tissue scale, especially in complex organs like the brain, STitch3D have showed its ability to draw comprehensive 3D maps and reveal 3D distributions of cell subtypes; On the organism scale, STitch3D successfully reconstructed the 3D whole organism atlas of the *Drosophila* embryo with identified 3D organ regions.

Recently, several computational methods are also developed for deciphering tissue structures by either identifying tissue regions or estimating cell-type distributions in ST data analysis. Specifically, methods including BayesSpace, SpaGCN and STAGATE are designed for handling the spatial domain detection task, while methods including RCTD, Cell2location, Tangram, DestVI and CARD are designed for solving the cell-type deconvolution task. While these existing computational methods have greatly facilitated the analysis of ST data, direct application of these methods did not sufficiently address all the critical 3D analysis tasks. STitch3D addresses these challenges and offers advantages over existing methods for 3D reconstruction by innovating on the data model.

First, building the 3D tissue model requires incorporation of multiple tissue slices collected along the *z*-axis of the tissue. However, most existing methods only focus on analyzing one single tissue slice, which prevents their effective application to the 3D construction problem. Different from existing methods, STitch3D enables joint modeling of gene expression levels and spatial locations from multiple tissue slices, where information across slices is effectively integrated for revealing 3D structure of tissues. By effectively designing the shared latent space and accounting for slice-specific and gene-specific effects, STitch3D is able to preserve biologically meaningful information that varies along the *z*-axis, while also removing batch effects across slices. Additionally, it has been shown by previous studies that integrating more data improves statistical power in single-cell data analysis [40, 41, 42, 43]. Here, as a large amount of information is shared across slices, such as similar gene expression levels in adjacent slices, STitch3D’s joint analysis of multiple slices also improves accuracy. Through comprehensive real data studies, we also demonstrated that incorporating more slices and borrowing information across them indeed helps STitch3D to gain higher accuracy in both 3D spatial domain detection and 3D cell-type deconvolution.

Second, existing methods are designed to perform either the spatial domain detection task or the cell-type deconvolution task. However, analysis results for characterizing complex tissue structures from the two tasks are inherently connected. Specifically, cell-type compositions of spots often change smoothly within a biologically meaningful spatial region, while they are less similar across distinct spatial regions. Hence, it is important to leverage information shared by the two tasks for 3D reconstruction of tissues. Different from existing methods, STitch3D addresses both the spatial domain detection task and the cell-type deconvolution task simultaneously in a unified framework, facilitating comprehensive characterization of tissue structures. With the design of simultaneously addressing the two 3D tasks, in STitch3D, cell-type transcriptomic profiles from single-cell reference datasets not only help deconvolve RNA expressions of spots, but also facilitate learning latent representations used for the spatial domain detection task. Since STitch3D links latent representations with cell-type proportions of spots using a shallow network, the cell-type compositions serve as augmented information for identifying biological interpretable 3D regions. Compared to existing spatial domain detection methods which assign domain labels in an unsupervised manner, STitch3D introduces additional signals for supervision.

In this study, we have focused on using scRNA-seq expression reference and spatially resolved gene expression measurements with spatial location information to reconstruct 3D tissue structures, and demonstrated that, by effectively integrating all related information, STitch3D has remarkable advantages over many existing methods. Effective extraction and utilization of H&E-stained histological image information, combined with our existing approach could further improve interpretability of results, and is left to future work.

3D reconstruction of tissue structures is evidently a very meaningful task in ST studies to accelerate biological discoveries. As more ST datasets containing multiple tissue slices are generated, the demand for using such datasets to assemble 3D panoramic views of tissues will grow rapidly. STitch3D caters to this demand for tools to reconstruct 3D tissue structures with multiple ST slices, and will play an essential role in ST data analysis.

## Methods

### The model of STitch3D

STitch3D first incorporates the iterative closest point (ICP) algorithm [44] or PASTE [10] algorithms to align multiple slices by registration of spatial spots. Details of the alignment step are presented in Supplementary section 3. After the alignment of multiple slices, STitch3D assembles the 2D locations of spots along a newly introduced *z*-axis, which depicts the distances between pairs of slices, to build 3D spatial locations of spots. With the 3D spatial locations of spots, STitch3D constructs a global 3D neighborhood graph for spots in all slices. By default, two spots are connected with an edge in the global 3D graph, if their distance is less than 1.1 times of the distance between two nearest spots in one slice.

Next, we present STitch3D model to perform 3D spatial domain detection and 3D cell type deconvolution. STitch3D performs 3D analyses by integrating information from all slices of a ST dataset and a scRNA-seq reference dataset. In the ST data, we use *s* = 1,2, ⋯, *S* as the index of tissue slices. For slice *s*, we denote its number of spots as *N_s_*, and the observed gene expression count matrix as 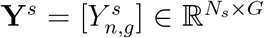, with index *n* = 1,2, ⋯, *N_s_* for spots and *g* = 1, 2, ⋯, *G* for genes. In the annotated scRNA-seq references, we denote the obtained cell type-specific gene expression profile as matrix 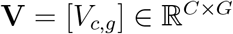, where *c* =1, 2, ⋯, *C* is the index of cell types. In this matrix, the row vector 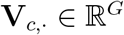, which satisfies 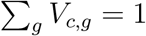, characterizes mean expression signature of cell type *c*.

To integrate both gene expression levels and spatial locations across all tissue slices for 3D analyses, STitch3D first encodes gene expression information of spots into a shared latent space, with a neural network incorporating our constructed 3D spatial locations. For clarity, we concatenate the index for spots across slices *s* = 1, 2, ⋯, *S* to introduce a new global index *i* = 1,2, ⋯, *N*, for spots in all slices where *N* = *N*_1_ + *N*_2_ + ⋯ + *N_S_*. We denote 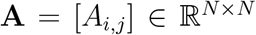 as the global adjacency matrix specifying the 3D global graph, where *A_i,j_* = 1 if there exists an edge between spot *i* and spot *j*, i.e., spot *i* and spot *j* are 3D neighboring spots, and *A_i,j_* = 0 otherwise. Let 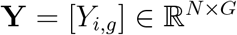 be the concatenated gene expression count matrix across all slices. Specifically, the latent representations of spots are obtained by:

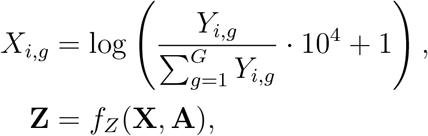

where we perform normalization and log-transformation on count matrix **Y** for network stability, 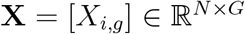 is encoded to 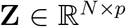 through a graph attention network *f_Z_*(·), and *p* is the dimensionality of the shared latent space. Clearly, the latent representation **Z** encodes information from both gene expression **X** and spatial neighbors given in graph **A**.

STitch3D then generates cell type proportions based on latent representations of spots. We denote the proportion of cell type *c* in spot *i* as *β_i,c_*. Let 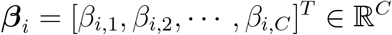 with 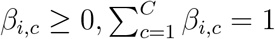 represent cell type proportions for spot *i*, and 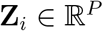 be the obtained latent representation for spot *i*, where *i* = 1,2, ⋯, *N*. We assume that cell type proportions ***β**_i_* can be related to latent representation **Z**_*i*_ through a neural network *f_β_*(·) as

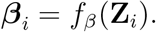

Of note, with graph attention network *f_Z_*(·), the obtained latent codes **Z** are aware of 3D global spatial information. Consequently, as ***β**_i_* is generated from **Z**_*i*_, spatial location information is also incorporated into the estimated cell type proportions *β_i_*.

To account for technical effects between ST data and scRNA-seq data, as well as batch effects among multiple ST slices, STitch3D further introduces two effects 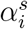 and 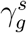 to its model, where 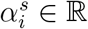 represents slice- and spot-specific effects, and 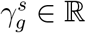 represents slice- and gene-specific effects. Combining cell type-specific gene expression profile matrix **V** = [*V_c,g_*], cell type proportions ***β**_i_* = [*β_i,c_*], as well as the two effects 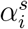 and 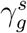, STitch3D is able to reconstruct the observed counts in ST data with the following model:

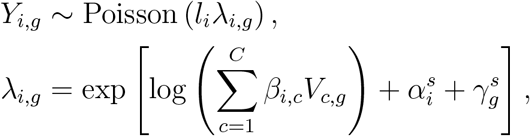

where *l_i_* is the observed total transcript count in spot *i*. Specifically, we generate slice- and spot-specific effects 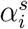 from the shared latent space by using slice-specific neural network:

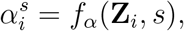

where *s* is the slice label. We model 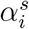 in such a way, because we assume slice- and spot-specific effects 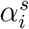 are related to both gene expression information which is distilled in **Z**_*i*_, and slice label information contained in *s*. We model slice- and gene-specific effects 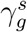 as trainable parameters. By accounting for unwanted variations in 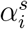 and 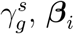 and **Z**_*i*_ can better capture biologically meaningful variation. Based on this model, the loss function is given by

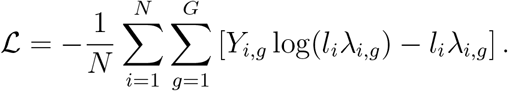

In our STitch3D model, we consider a regularizer to encourage the preservation of biological variations across multiple slices. Specifically, we design the regularizer as

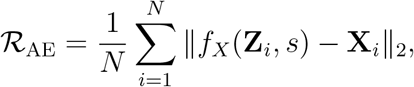

where slice-specific network *f_X_*(·, *s*) is used to reconstruct **X**_*i*_ based on **Z**_*i*_ and slice label *s*. The two networks, *f_Z_*(·) and *f_X_*(·, *s*), together form an auto-encoder structure between **X** and **Z**. In regularizer 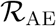, the slice-specific network *f_X_*(·, *s*) is designed to account for batch effects among slices, therefore encouraging slice-shared network *f_Z_*(·) to distill biological information in **Z**. The final objective function in STitch3D is given by 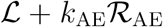, where *k*_AE_ is the coefficient for regularizer 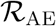 and pre-fixed at 0.1.

With above designs, STitch3D applies stochastic gradient descent to update trainable parameters in its model. After training, STitch3D outputs latent representations **Z**_*i*_ and cell type proportions ***β**_i_*. The latent representations are utilized for the 3D spatial domain identification task with community detection algorithms, such the Louvain algorithm [45] and Gaussian mixture model-based clustering algorithm. Combining with obtained 3D spatial locations, such identified clusters reveal 3D spatial domains across all slices. Meanwhile, cell type proportions ***β**_i_* show the 3D spatial distributions of different cell types, providing a comprehensive view for ST studies in a higher resolution.

### Selection of genes and construction of cell type-specific gene expression profile matrix

STitch3D selects informative genes based on the scRNA-seq reference data. Specifically, for each cell type in the annotated scRNA-seq data, STitch3D uses *t*-test to find its top *K* marker genes with default setting *K* = 500. Then the top marker genes for all cell types are concatenated to get a full list of informative genes that are used by STitch3D.

With *G* selected genes from *C* cell types, STitch3D constructs the cell type-specific gene expression profile matrix 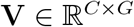. First, STitch3D normalizes gene expression measurements for each cell in the reference scRNA-seq dataset, such that the sum of gene expressions of each cell equals to 1. Then, the row vector **V**_*c*_,., the mean expression signature of cell type c, is computed by averaging all normalized expressions of cells with cell type c. If different batches are included in the reference scRNA-seq dataset, then STitch3D computes a cell type-specific gene expression profile matrix for each batch, and takes average of them to get the overall cell type-specific gene expression profile matrix **V**.

### Network structures in STitch3D

For encoder network *f_Z_*(·) which extracts latent representations of spots in the shared latent space, STitch3D adopts a graph attention network to incorporate the constructed 3D spatial graph. The graph attention network contains a graph attention layer and a dense layer. Inputs to the graph attention layer in *f_Z_*(·) are global adjacency matrix **A**, and normalized and log-transformed gene expression **X**. The dimensionality of **X** is the number of selected highly variable genes *G* that varies from dataset to dataset. The dimensionality of the output to graph attention layer is pre-fixed at 512. Then the dense layer in *f_Z_*(·) takes the 512-dimensional hidden vectors as input and outputs 128-dimensional latent representations **Z**. For network *f_β_*(·) which produces cell type proportions, STitch3D adopts a one-layer dense network to map 128-dimensional latent representations **Z** to *C*-dimensional outputs, where *C* is the number of cell types, with the softmax activation. STitch3D adopts a one-layer dense network for *f_α_*(·, *s*) which takes latent representations **Z** and slice labels as inputs, and generates slice- and spot-specific effects 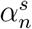. For the slice-specific decoder network *f_X_*(·, *s*) in the regularizer 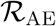, a graph attention network with hidden dimensionality 512 is used. Specifically, inputs to the graph attention layer are global adjacency matrix **A**, latent representations **Z**, and slice labels. The dimensionality of the output to graph attention layer is 512. Then a dense layer is used, which takes the 512-dimensional hidden vectors as input, and reconstructs gene expression matrix.

To show how to leverage spatial location information in STitch3D, we describe details of graph attention layers used in encoder network *f_Z_*(·) and decoder network *f_X_*(·, *s*). Let embedding 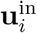 of spot *i* be the input to the graph attention layer. The graph attention layer updates spot embedding by using global adjacency matrix 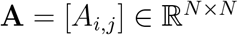 of the constructed 3D global graph and the graph attention mechanism:

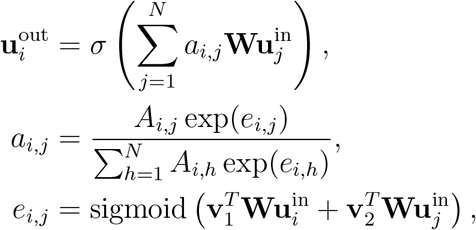

where *σ*(·) is the activation function, **W** is the network parameter in the graph attention layer, **v**_1_ and **v**_2_ are the parameters related to the graph attention mechanism. Parameters **v**_1_ and **v**_2_ are used to learn edge weights *a_i,j_* between 3D neighborhood spots during training, helping to borrow information from neighborhood spots adaptively for 3D analyses. For encoder network *f_Z_*(·), 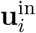 is gene expression-level data **X**_*i*_, and 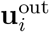 is a hidden vector with dimensionality 512. For decoder network *f_X_*(·, *s*), 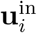 is latent representation **Z**_*i*_, and 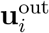 is a hidden vector with dimensionality 512.

### Model training details

STitch3D utilizes Adamax, which is a variant of Adam algorithm [46], for stochastic optimization in its model training. By default, the number of optimization steps in STitch3D is set to 20,000 with learning rate *lr* = 0.002, coefficients for computing running averages *β*_1_ = 0.9, *β*_2_ = 0.999, and weight decay parameter λ = 0. As demonstrated in Supplementary Fig. 23, STitch3D’s training process guarantees the convergence of objective functions for large datasets such as the mouse brain dataset [6] with 17, 086 spots. We ran all experiments using one single GPU, and the timings of training processes for these ST datasets are presented in Supplementary Table 2.

### Denoising and imputation of gene expression levels

STitch3D’s cell type deconvolution results also enable denoising of expression levels of low-quality genes and imputatiuon of unmeasured gene expressions in ST datasets. For the genes of interest to be denoised or imputed, STitch3D first calculates a cell type-specific gene expression profile vector 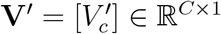 based on the scRNA-seq reference, in the same way how the cell type-specific gene expression profile matrix **V** of highly variable genes is obtained. Then, using STitch3D’s estimated cell type proportion in each spot *β_i_*, the denoised or imputed expression levels of the genes of interest are given by 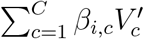.

## Data availability

All data used in this work are publicly available through online sources.

- Human dorsolateral prefrontal cortex dataset profiled by Visium platform [7] (http://spatial.libd.org/spatialLIBD/).
- Human dorsolateral prefrontal cortex dataset profiled by 10x Genomics Chromium platform [22] (GSE144136).
- Mouse cortex dataset profiled by seqFISH+ [24] (https://github.com/CaiGroup/seqFISH-PLUS).
- Mouse primary visual cortex dataset profiled by SMART-seq [27] (https://portal.brain-map.org/atlases-and-data/rnaseq/mouse-v1-and-alm-smart-seq).
- Mouse visual cortex dataset profiled by STARmap [25] (https://www.starmapresources.com/data).
- Mouse hypothalamic preoptic dataset profiled by MERFISH [26] ([Dryad](https://datadryad.org/stash/dataset/doi:10.5061/dryad.8t8s248)).
- Mouse hypothalamic preoptic dataset profiled by Illumina NextSeq 500 [26](GSE113576).
- Mouse whole brain dataset profiled by ST platform [6] (GSE147747).
- Mouse brain dataset profiled by 10x Genomics Chromium platform [6] (E-MTAB-11115).
- Human embryonic heart dataset profiled by ST platform [8] (https://data.mendeley.com/datasets/dgnysc3zn5/1).
- Human embryonic heart dataset profiled by 10x Genomics Chromium platform [8] (https://data.mendeley.com/datasets/mbvhhf8m62/2).
- *Drosophila* embryo dataset profiled by Stereo-seq [9] (https://db.cngb.org/stomics/flysta3d/download.html).
- *Drosophila* embryo dataset profiled by sci-RNA-seq [35] (GSE190149).

## Code availability

STitch3D software is available at https://github.com/YangLabHKUSl7STitch3D.

## Acknowledgements

We acknowledge grants as follows: Hong Kong Research Grant Council Grants 16307818, 16301419, 16308120, 16307221, Hong Kong Innovation and Technology Fund Grant PRP/029/19FX, Hong Kong University of Science and Technology Startup Grants R9405, Z0428 from the Big Data Institute, Guangdong-Hong Kong-Macao Joint Laboratory Grant no. 2020B1212030001 and the RGC Collaborative Research Fund Grant C6021-19EF to C.Y.; Hong Kong Research Grant Council Grant 16209820, Lo Ka Chung Foundation through the Hong Kong Epigenomics Project, Chau Hoi Shuen Foundation, the SpatioTemporal Omics Consortium (STOC) and the STOmics Grant Program to A.R.W.; Hong Kong Research Grant Council Grant 16103620, the Shenzhen Science and Technology Innovation Commission JCYJ20180223181229868 and JCYJ20200109140201722 to Y.Y.

