## Supplementary Information for "Construction of a 3D whole organism spatial atlas by joint modeling of multiple slices"

#### Contents

**Supplementary Section 1: Settings and results in the three scenarios of the simulation study.** **2**

**Supplementary Section 2: Evaluation metrics used in the simulation studies.** **4**

**Supplementary Section 3: Alignment of multiple slices and construction of 3D global graph.** **5**

*These authors contributed to this work equally.

**Supplementary Figures** **7**

**Supplementary Tables** **20**

### Supplementary Section 1: Settings and results in the three scenarios of the simulation study.

We followed a recent study [1] to compare STitch3D and other cell-type deconvolution methods, including RCTD [2], Cell2location [3], Tangram [4], DestVI [5] and CARD [6], based on simulations. These methods were compared in three different scenarios. In each scenario, simulated spatial transcriptomics (ST) slices were created by gridding cells with spatial locations into multi-cell spots. To have a fair comparison, the output of each method was normalized such that the sum of all cell-type proportions in each spot equals to one. To quantitatively compare estimated cell-type proportions with the ground truth in each spot, we evaluated performance metrics including the Pearson correlation coefficient (PCC), structural similarity index (SSIM) [7], root mean square error (RMSE) and Jensen–Shannon distance (JS). An accuracy score of each method was computed based on the metrics. Details of performance metrics can be found in Supplementary Section 2.

In the first scenario, we compared all methods in single-slice setting. We used a mouse cortex seqFISH+ dataset [8] to create a single simulated ST slice by gridding cells into spots (Supplementary Fig. 3**a**). With a mouse primary visual cortex single-cell dataset profiled by SMART-seq [9] serving as the cell-type expression reference, we applied all methods to this dataset for the cell-type deconvolution task. In this experiment, STitch3D correctly recovered spatial distributions of different cell types, such as the excitatory neurons in layers L5-L6 (Supplementary Fig. 3**b**). Additionally, STitch3D achieved an overall accuracy score equal to exactly one in the quantitative evaluation result (Supplementary Fig. 3**c**), indicating that it outperformed other compared methods in terms of all four metrics. Besides cell-type deconvolution, with this dataset, we also used STitch3D for the gene imputation task. Three genes including *Cplx1*, *Igsf21* and *Rprm* were manually removed from the simulated ST slice. Then STitch3D was applied to predict the expressions of these three genes in z-scores. Compared to the ground truth gene expressions, STitch3D showed reliable gene imputation performance. For example, it correctly recovered the gene expression pattern of *Cplx1* (Supplementary Fig. 3**d**). Among compared methods, Tangram [4] is also capable of predicting unmeasured genes in the ST dataset, so we compared STitch3D with Tangram in this task. We used PCC, SSIM, RMSE and JS to quantify the similarity of predicted gene expressions and ground truth gene expressions in z-scores. As shown in the result (Supplementary Fig. 3**e**), for all three genes, STitch3D consistently achieved better scores, demonstrating its high accuracy in gene imputation.

We then compared all methods in the second scenario. Multiple slices were created from a single mouse visual cortex STARmap slice [10] as technical replicates, by adding independent standard Gaussian noises to log-transformed gene expressions (Supplementary Fig. 4**a**). The single-cell expression reference dataset in the first scenario was also used for this scenario. We compared all methods in single-slice setting. As Cell2location is also able to jointly deal with multiple slices, we compared it with STitch3D in multi-slice setting. According to the results, STitch3D is among the best two methods in single-slice experiments (Supplementary Fig. 4**c**). When more slices were incorporated, STitch3D showed stably improved accuracy scores (Supplementary Fig. 4**c**). Its deconvolution result also became less noisy when more slices were included (Supplementary Fig. 4**b**). However, Cell2location showed less satisfactory performance in multi-slice setting, with decreasing accuracy scores as more slices were used (Supplementary Fig. 4**c**).

In the third scenario, a more realistic multi-slice simulation setting was introduced. Instead of simulating multiple technological replicates from one single slice, we created multiple slices from adjacent mouse hypothalamic preoptic region MERFISH slices [11] (Supplementary Fig. 5**a**). The scRNA-seq dataset provided in the same study was used as the single-cell reference. In single-slice experiments, all methods were applied to the center slice (Bregma $-0.24$) among the three adjacent slices. Among compared methods, STitch3D showed the best performance in the cell-type deconvolution task for a single slice (Supplementary Fig. 5**b**, **c**). We then applied STitch3D and Cell2location to jointly deal with the three adjacent slices. The result indicated that STitch3D can effectively borrow information across slices. We also repeated this analysis to the other two sets of three adjacent slices, with center slices at Bregma $-0.19$ and $-0.14$ respectively. The experiments showed consistent results (Supplementary Fig. 5**d**, **f**), demonstrating STitch3D's ability to joint modeling of multiple slices in the cell-type deconvolution task.

### Supplementary Section 2: Evaluation metrics used in the simulation studies.

We compared cell-type deconvolution methods with metrics including the Pearson correlation coefficient (PCC), structural similarity index (SSIM) [7], root mean square error (RMSE) and Jensen–Shannon distance (JS), following the benchmarking study [1].

Let $n = 1, 2, \cdots, N$ be index for spots and $c = 1, 2, \cdots, C$ be index for cell types. Let $\boldsymbol{\beta}_n = [\beta_{n,1}, \beta_{n,2}, \cdots, \beta_{n,C}]^T$ be estimated cell-type proportions in the $n$-th spot, and $\tilde{\boldsymbol{\beta}}_n = [\tilde{\beta}_{n,1}, \tilde{\beta}_{n,2}, \cdots, \tilde{\beta}_{n,C}]^T$ be ground truth cell-type proportions in the $n$-th spot. We only use shared cell types between the ground truth and the single-cell reference for evaluation of metrics.

PCC of the $n$-th spot is calculated as

$$\mathrm{PCC}_n = \frac{\sum_{c=1}^{C}(\beta_{n,c} - \mu_n)(\tilde{\beta}_{n,c} - \tilde{\mu}_n)}{\sigma_n \tilde{\sigma}_n},$$

where

$$\mu_n = \frac{1}{C}\sum_{c=1}^{C}\beta_{n,c},\ \tilde{\mu}_n = \frac{1}{C}\sum_{c=1}^{C}\tilde{\beta}_{n,c},\ \sigma_n = \sqrt{\sum_{c=1}^{C}(\beta_{n,c} - \mu_n)^2},\ \tilde{\sigma}_n = \sqrt{\sum_{c=1}^{C}(\tilde{\beta}_{n,c} - \tilde{\mu}_n)^2}.$$

A higher PCC indicates a better performance.

To calculate SSIM of the $n$-th spot, $\boldsymbol{\beta}_n$ and $\tilde{\boldsymbol{\beta}}_n$ are first rescaled by

$$\boldsymbol{\beta}'_n = \boldsymbol{\beta}_n / \max\{\beta_{n,1}, \beta_{n,2}, \cdots, \beta_{n,C}\},\ \tilde{\boldsymbol{\beta}}'_n = \tilde{\boldsymbol{\beta}}_n / \max\{\tilde{\beta}_{n,1}, \tilde{\beta}_{n,2}, \cdots, \tilde{\beta}_{n,C}\},$$

such that their largest entries equal to one. Then SSIM of the $n$-th spot is calculated as

$$\mathrm{SSIM}_n = \frac{2\mu'_n \tilde{\mu}'_n + C_1^2}{(\mu'_n)^2 + (\tilde{\mu}'_n)^2 + C_1^2} \cdot \frac{2\mathrm{Cov}(\boldsymbol{\beta}'_n, \tilde{\boldsymbol{\beta}}_n) + C_2^2}{(\sigma'_n)^2 + (\tilde{\sigma}'_n)^2 + C_2^2},$$

where $C_1 = 0.01$, $C_2 = 0.03$,

$$\mu'_n = \frac{1}{C}\sum_{c=1}^{C}\beta'_{n,c},\ \tilde{\mu}'_n = \frac{1}{C}\sum_{c=1}^{C}\tilde{\beta}'_{n,c},\ \sigma'_n = \sqrt{\sum_{c=1}^{C}(\beta'_{n,c} - \mu'_n)^2},\ \tilde{\sigma}'_n = \sqrt{\sum_{c=1}^{C}(\tilde{\beta}'_{n,c} - \tilde{\mu}'_n)^2}.$$

A higher SSIM indicates a better performance.

RMSE of the $n$-th spot is calculated as

$$\mathrm{RMSE}_m = \sqrt{\frac{1}{C}\sum_{c=1}^{C}(\tilde{\beta}_{n,c} - \beta_{n,c})^2},$$

where a lower RMSE indicates a better performance.

JS computes the Jensen–Shannon distance between two probability vectors. JS of the $n$-th spot is given by the square root of the Jensen–Shannon divergence:

$$\mathrm{JS}_n = \sqrt{D_{\mathrm{JS}}\left[\boldsymbol{\beta}_n/(C\mu_n) \| \tilde{\boldsymbol{\beta}}_n/(C\tilde{\mu}_n)\right]},$$

where

$$D_{\mathrm{JS}}\left[\mathbf{p}\|\mathbf{q}\right] = \frac{1}{2}D_{\mathrm{KL}}\left[\mathbf{p}\|(\mathbf{p}+\mathbf{q})/2\right] + \frac{1}{2}D_{\mathrm{KL}}\left[\mathbf{q}\|(\mathbf{p}+\mathbf{q})/2\right],$$

and $D_{\mathrm{KL}}\left[\mathbf{a}\|\mathbf{b}\right] = \sum_{c=1}^{C} a_c \log \frac{a_c}{b_c}$ is the Kullback-Leibler divergence between $C$-dimensional probability arrays $\mathbf{a}$ and $\mathbf{b}$. A lower JS indicates a better performance.

After calculating these metrics, four sets of ranks among the $M$ methods were obtained, i.e., $\mathrm{Rank}_{PCC}$, $\mathrm{Rank}_{SSIM}$, $\mathrm{Rank}_{RMSE}$ and $\mathrm{Rank}_{JS}$. Each set of ranks ranges from 1 to $M$, such that a higher rank value means a better performance in terms of the corresponding metric. The overall accuracy score of a method was computed by leveraging its ranks with respect to the four metrics:

$$\text{Accuracy score} = \frac{1}{4M}(\mathrm{Rank}_{PCC} + \mathrm{Rank}_{SSIM} + \mathrm{Rank}_{RMSE} + \mathrm{Rank}_{JS}).$$

If the accuracy score of a method equals to one, it means that this method outperforms the others in all four metrics.

Similar to the benckmarking in terms of cell-type deconvolution task, to evaluate the performance of gene imputation, the above four metrics can be calculated by replacing estimated and ground truth cell-type proportions with predicted and ground truth gene expressions respectively.

### Supplementary Section 3: Alignment of multiple slices and construction of 3D global graph.

STitch3D aligns multiple slices by registration of spatial spots. By default, STitch3D adopts the iterative closest point (ICP) algorithm [12] to align slices pairwise. Before alignment, STitch3D detects edge spots in each slice based on the numbers of neighborhoods. For example, for datasets acquired with the Visium platform, a non-edge spot should have six neighborhood spots. For robustness, we define an edge spot with the condition that its number of neighborhood spots is less than five and more than one. In datasets profiled with the ST platform, a non-edge spot should have eight neighborhood spots. Correspondingly, an edge spot is identified if it has less than seven and more than one neighborhood spots. Next, STitch3D uses the ICP algorithm to pairwise align edge spots from two slices, assuming that edge spots are usually located on the boundary of a tissue. Let $P = \{\mathbf{p}_1, \mathbf{p}_2, \cdots, \mathbf{p}_m\}$ be the set of locations of edge spots in the source slice, and $Q = \{\mathbf{q}_1, \mathbf{q}_2, \cdots, \mathbf{q}_k\}$ be the set of locations of edge spots in the target slice, where $\mathbf{p}_i, \mathbf{q}_i \in \mathbb{R}^2$. The ICP algorithm aligns spots by iteratively performing the following steps until convergence:

1. For each point in the source point cloud $P$, find its closest point in the target point cloud $Q$ to form a new set $Q'$. Then, $Q' = \{\mathbf{q}'_1, \mathbf{q}'_2, \cdots, \mathbf{q}'_m\}$ containing points in the target point cloud is obtained.

2. Find the transformation with the optimal rotation and translation by solving the least square problem:

$$\hat{\mathbf{R}}, \hat{\mathbf{v}} = \underset{\mathbf{R}\in\mathbb{R}^{2\times 2}, \mathbf{v}\in\mathbb{R}^2}{\operatorname{argmin}} \sum_{i=1}^{m} \left\| (\mathbf{R}\mathbf{p}_i + \mathbf{v}) - \mathbf{q}'_i \right\|^2,$$
$$\text{subject to} \quad \mathbf{R}^T\mathbf{R} = \mathbf{I}.$$

3. Apply the transformation $\hat{\mathbf{R}}\mathbf{p}_i + \hat{\mathbf{v}}$ to the source point cloud $P$.

For multiple slices, STitch3D performs pairwise alignment provided by the ICP algorithm sequencially. Except the ICP algorithm, STitch3D also incorporates a recently proposed method named PASTE, as described in its original paper [13], as an optional choice for the pairwise alignment task.

### Supplementary Figures

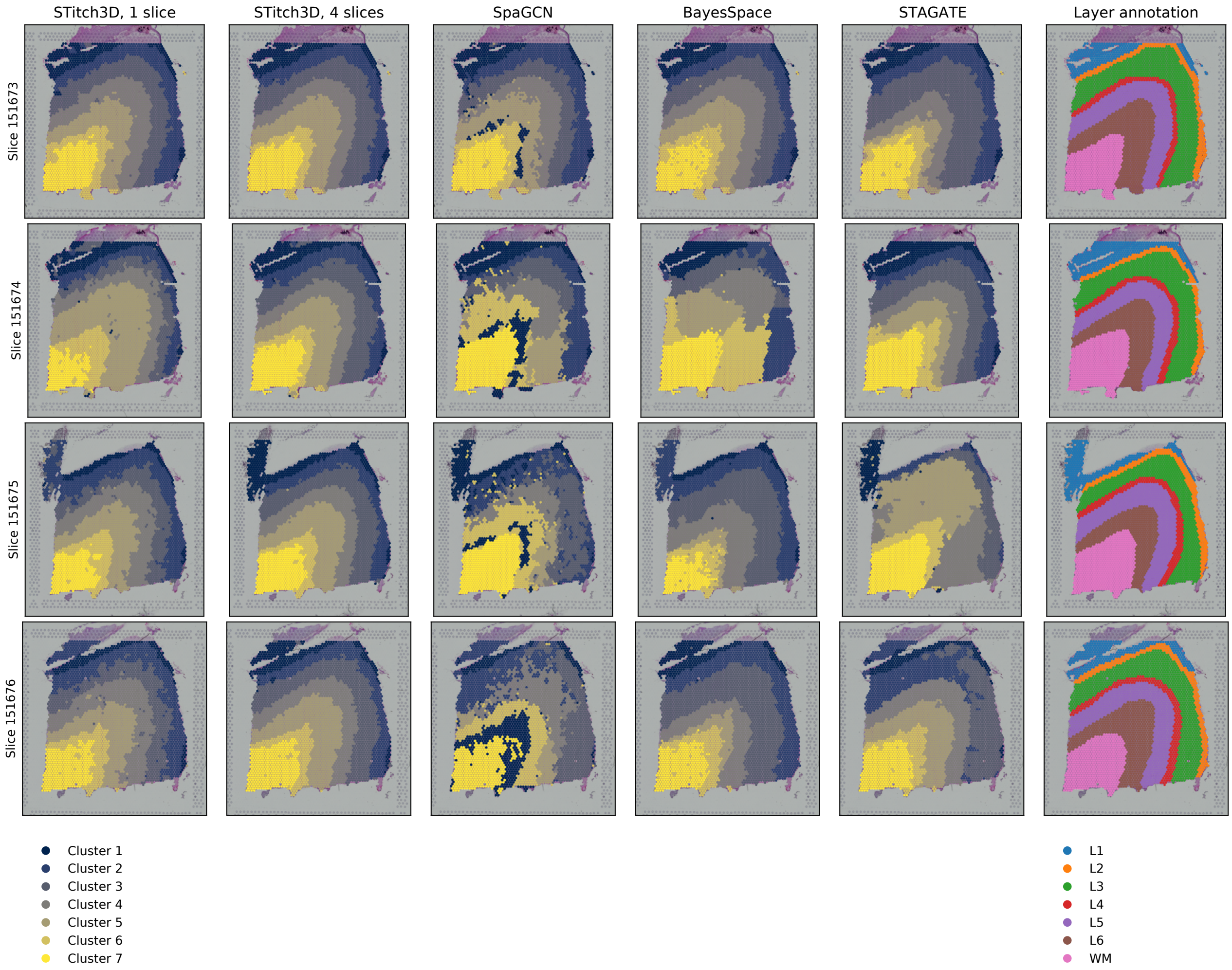


Supplementary Fig. 1: **Comparison of spatial domain detection methods based on the human dorsolateral prefrontal cortex (DLPFC) dataset [14].** We visualized spatial domain detection results of all compared methods, and ground truth layer annotation. STitch3D was used in both single-slice and multi-slice settings.

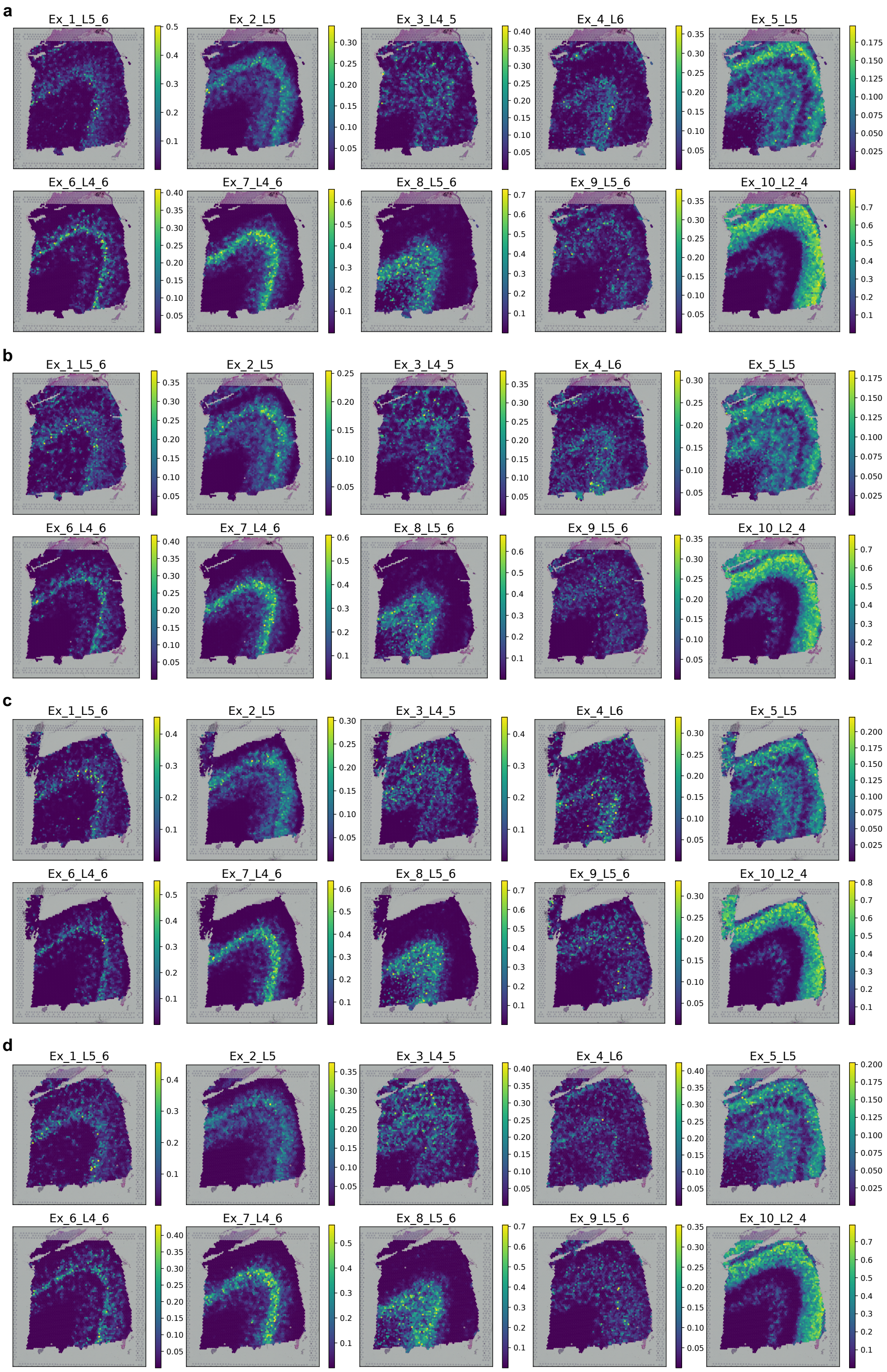


Supplementary Fig. 2: **STitch3D identified cell-type distributions in the DLPFC dataset.** STitch3D's cell-type deconvolution results in multi-slice setting on ten neuronal subtypes. **a**. Slice 151673. **b**. Slice 151674. **c**. Slice 151675. **d**. Slice 151676.

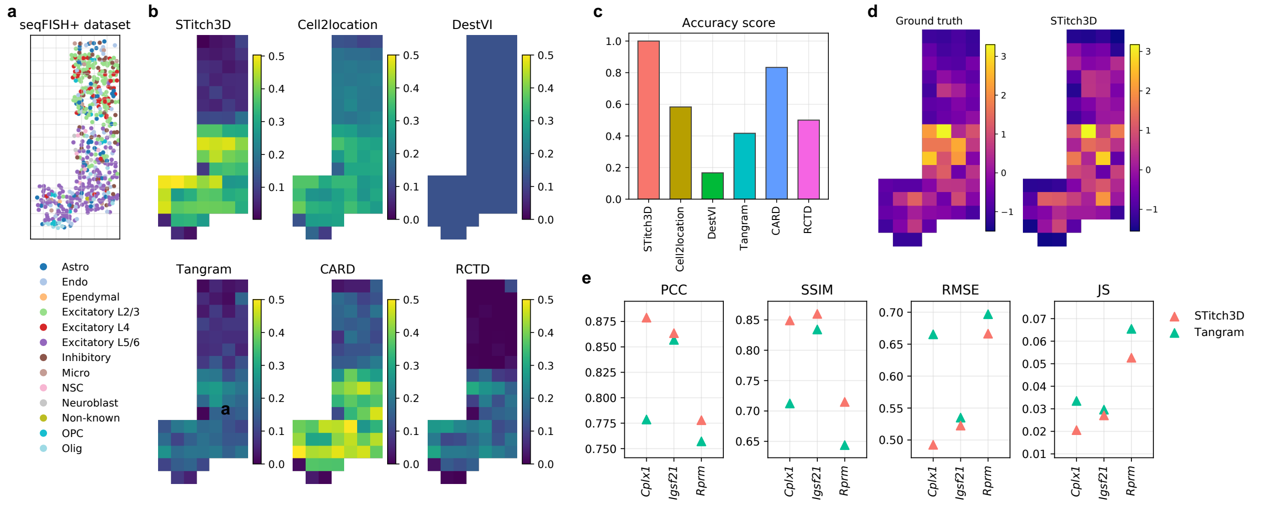


Supplementary Fig. 3: **Cell-type deconvolution results based on mouse cortex seqFISH+ data [8].** **a**. The seqFISH+ slice used for the simulation setting in the first scenario. **b**. Comparison of cell-type deconvolution results of excitatory L5/6 neuronal subtype. **c**. Overall accuracy scores of compared methods. **d**. Ground truth *Cplx1* expression (left) and STitch3D's imputation result (right) in z-scores. **e**. Quantitative comparison between STitch3D's and Tangram's imputation results of genes *Cplx1*, *Igsf21* and *Rprm*. Higher PCC, SSIM and lower RMSE, JS scores indicate better performance.

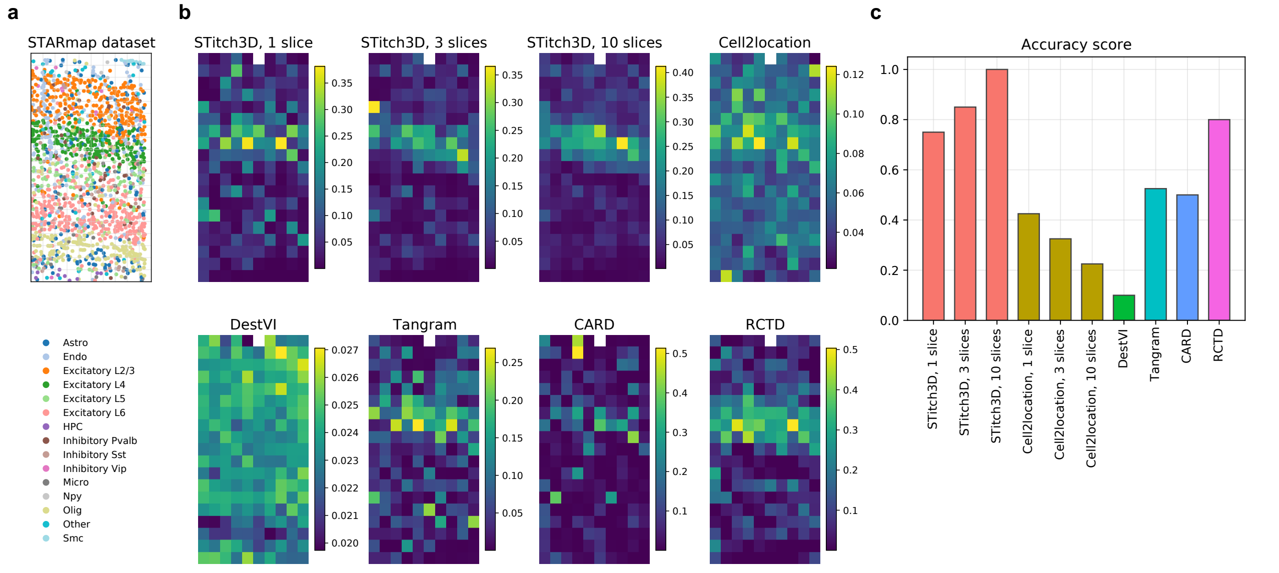


Supplementary Fig. 4: **Cell-type deconvolution results based on mouse visual cortex STARmap data [10].** **a**. The STARmap slice used for the simulation setting in the second scenario. **b**. Comparison of cell-type deconvolution results of excitatory L4 neuronal subtype. **c**. Overall accuracy scores of compared methods.

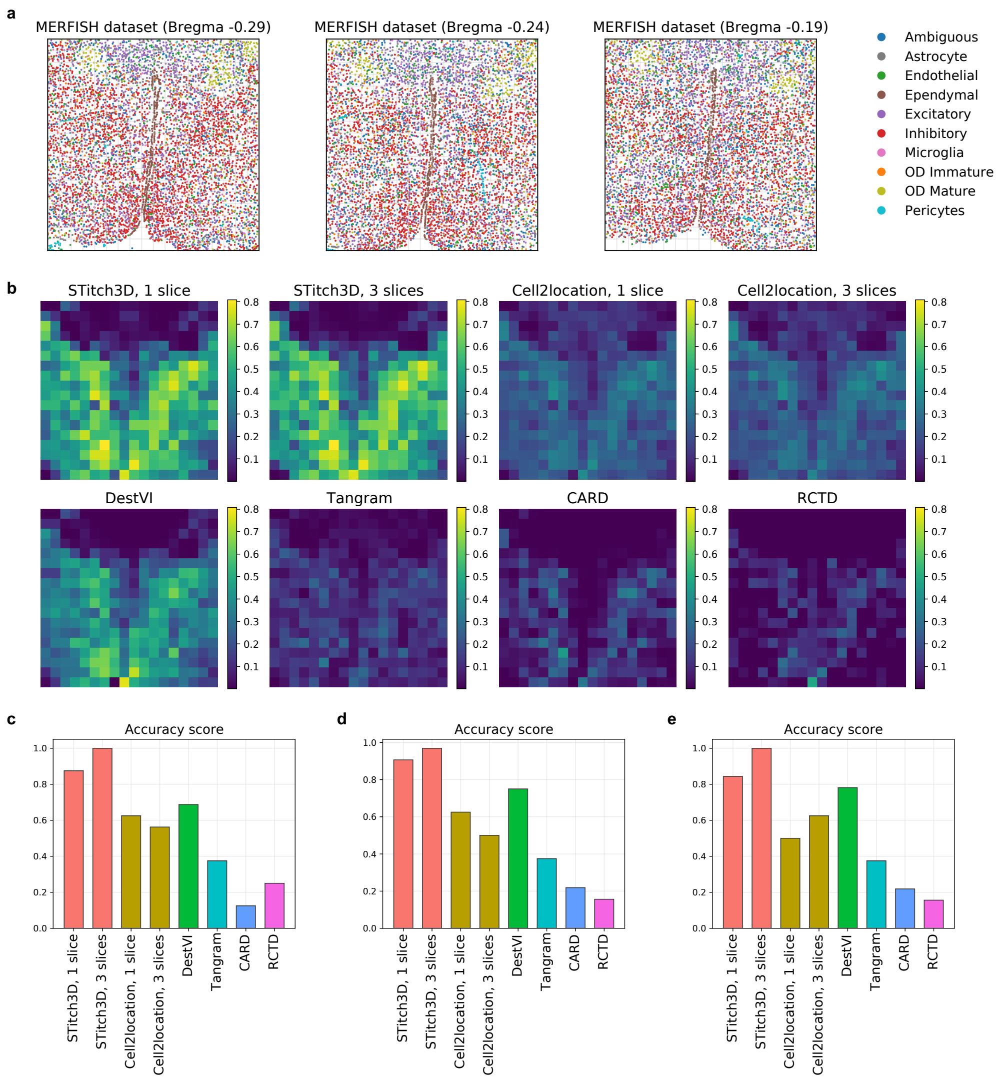


Supplementary Fig. 5: **Cell-type deconvolution results based on mouse hypothalamic preoptic region MERFISH data [11].** **a**. Three adjacent MERFISH slices at Bregma −0.29, −0.24 and −0.19 used for the simulation setting in the third scenario. **b**. Comparison of cell-type deconvolution results of inhibitory neurons for the center slice (Bregma −0.24). **c**. Overall accuracy scores of compared methods with three slices at Bregma −0.29, −0.24 and −0.19. Single-slice experiments were tested on the center slice at Bregma −0.24. **d**. Overall accuracy scores of compared methods with three slices at Bregma −0.24, −0.19 and −0.14. Single-slice experiments were tested on the center slice at Bregma −0.19. **c**. Overall accuracy scores of compared methods with three slices at Bregma −0.19, −0.14 and −0.09. Single-slice experiments were tested on the center slice at Bregma −0.14.

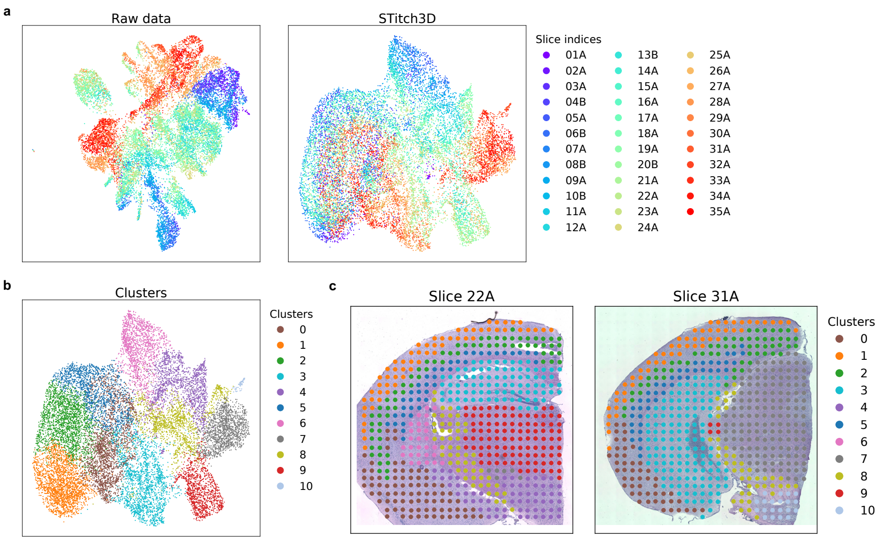


Supplementary Fig. 6: **STitch3D's spatial domain detection result for the adult mouse brain [15]. a**. UMAP plots [16] of raw data (left) and STitch3D's learned representation in the shared latent space (right), colored by slice indices. **b**. UMAP plot of STitch3D's learned representation colored by cluster labels. **c**. STitch3D's spatial domain detection result visualized on two slices.

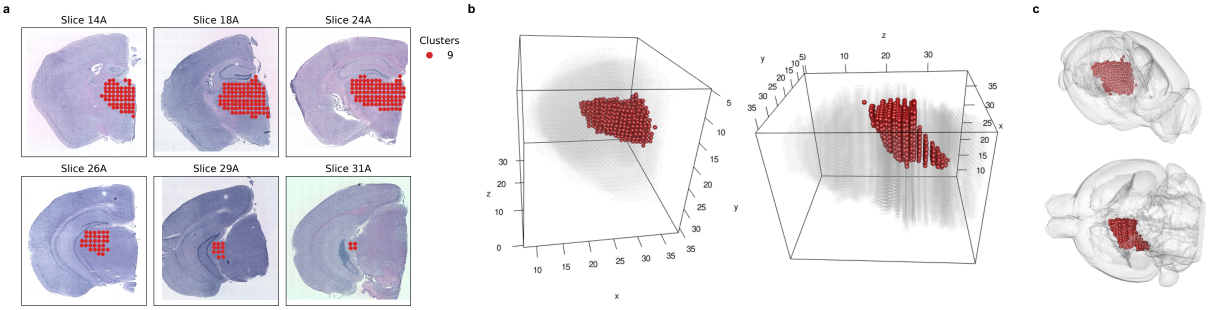


Supplementary Fig. 7: **Cluster 9 in STitch3D's spatial domain detection result for the adult mouse brain. a**. Cluster 9 visualized on six slices. **b**. 3D visualizations of cluster 9 in STitch3D's aligned 3D coordinates. **c**. 3D visualizations of cluster 9 in the Allen Mouse Brain Common Coordinate Framework (CCFv3) [17].

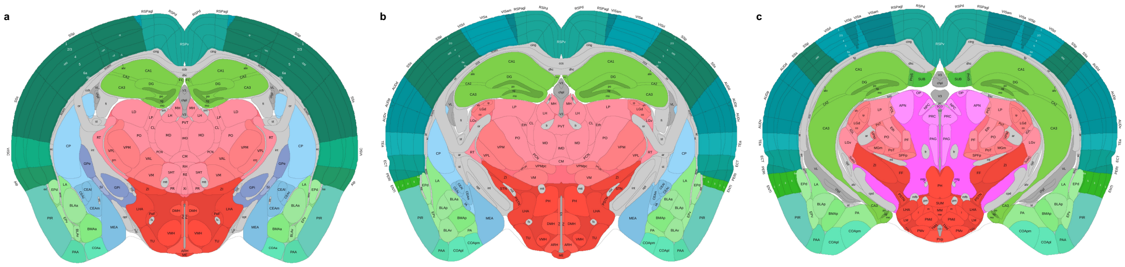


Supplementary Fig. 8: **Mouse brain coronal anatomical reference. a-c**. Three mouse brain coronal anatomical reference slices with 500 µm intervals between **a** and **b** and between **b** and **c**, provided by the Allen Reference Atlas – Mouse Brain [18], `http://atlas.brain-map.org/atlas?atlas=602630314`.

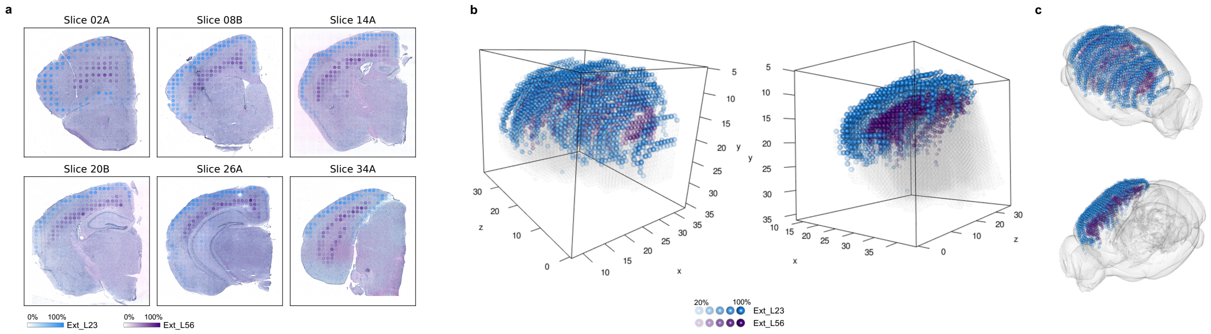


Supplementary Fig. 9: **Spatial distributions of Ext_L23 and Ext_L56 in STitch3D's cell-type deconvolution result for the adult mouse brain. a**. Estimated proportions of cell types Ext_L23 and Ext_L56 in STitch3D's cell-type deconvolution result visualized on 2D ST slices. A lower transparency indicates a higher proportion. **b**, **c**. 3D visualizations of proportions of cell types Ext_L23 and Ext_L56 in STitch3D's aligned 3D coordinates using ICP (**b**) and in CCFv3 (**c**), where spots with proportion values larger than 20% are shown.

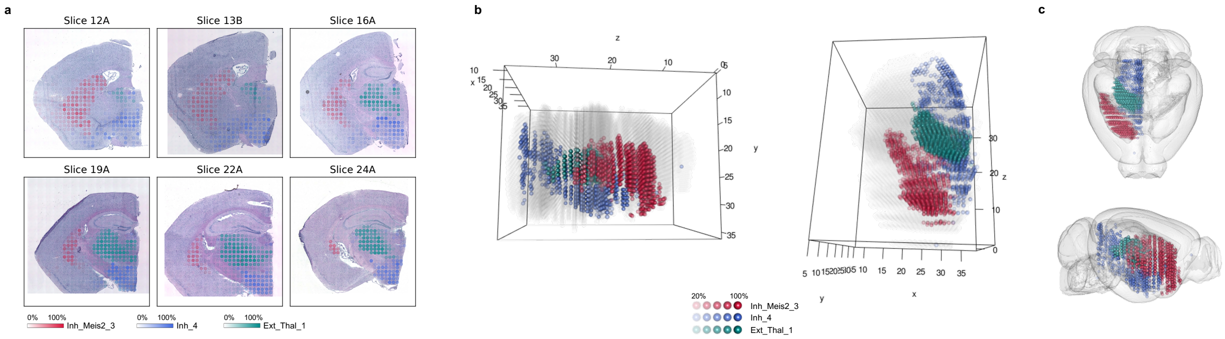


Supplementary Fig. 10: **Spatial distributions of Inh_Meis2_3, Inh_4 and Ext_Thal_1 in STitch3D's cell-type deconvolution result for the adult mouse brain.** **a**. Estimated proportions of cell types Inh_Meis2_3, Inh_4 and Ext_Thal_1 in STitch3D's cell-type deconvolution result visualized on 2D ST slices. A lower transparency indicates a higher proportion. **b**, **c**. 3D visualizations of proportions of cell types Inh_Meis2_3, Inh_4 and Ext_Thal_1 in STitch3D's aligned 3D coordinates using ICP (**b**) and in CCFv3 (**c**), where spots with proportion values larger than 20% are shown.

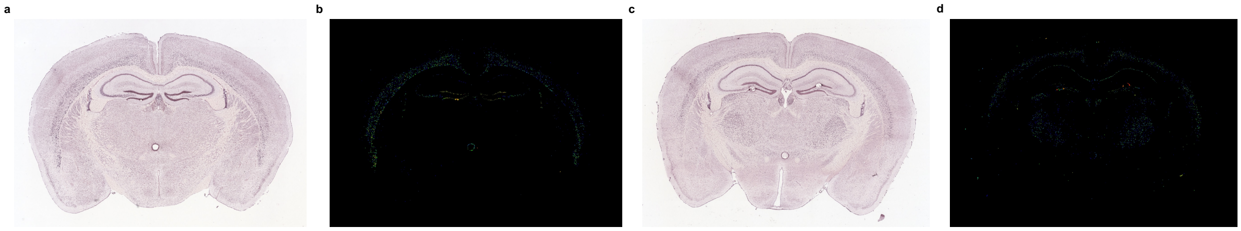


Supplementary Fig. 11: **Expression references of genes *Tle4* and *Rell1*.** **a**, **b**. *In situ* hybridization (ISH) image (**a**) and signal intensity heat map (**b**) of gene *Tle4* provided by the Allen Mouse Brain Atlas [19], `http://mouse.brain-map.org/experiment/show/73521809`. **c**, **d**. ISH image (**c**) and signal intensity heat map (**d**) of gene *Rell1* provided by the Allen Mouse Brain Atlas, `http://mouse.brain-map.org/experiment/show/72007746`.

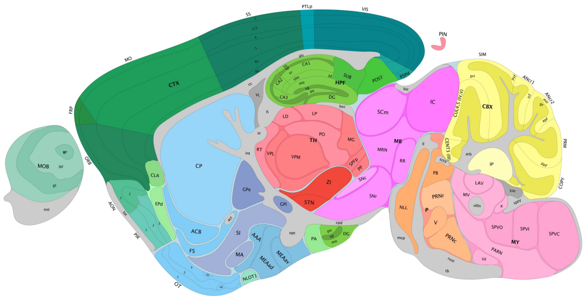

Supplementary Fig. 12: **Mouse brain sagittal anatomical reference.** A mouse brain sagittal anatomical reference slice provided by the Allen Reference Atlas – Mouse Brain [18], `http://atlas.brain-map.org/atlas?atlas=2&plate=100884129`.

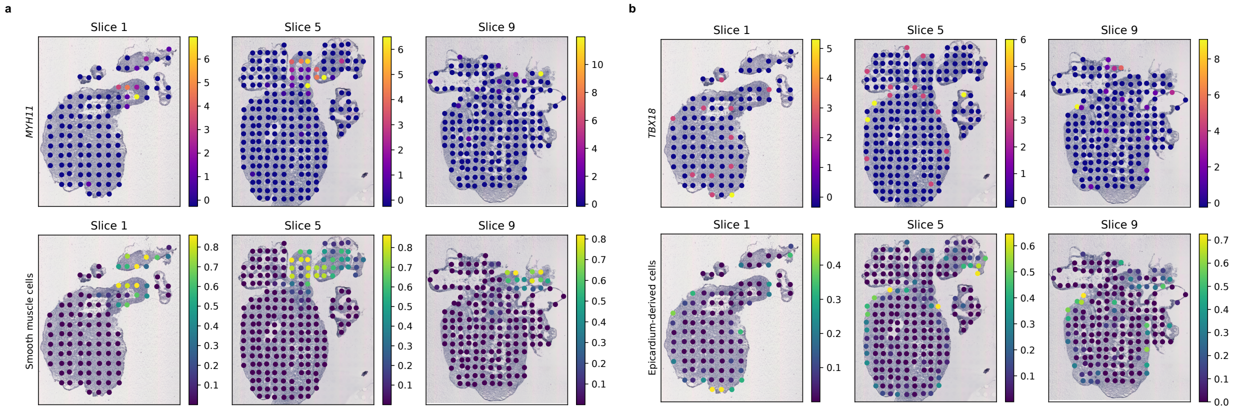


Supplementary Fig. 13: **Comparison between marker genes and STitch3D's cell-type deconvolution result for the 6.5 post-conception week (PCW) human heart. a**. Expression pattern of smooth muscle-related marker gene *MYH11* (the first row) and STitch3D's estimated cell-type proportions of smooth muscle cells (the second row) visualized on three ST slices. **b**. Expression pattern of epicardium-derived cell (EPDC) marker gene *TBX18* (the first row) and STitch3D's estimated cell-type proportions of EPDCs (the second row) visualized on three ST slices.

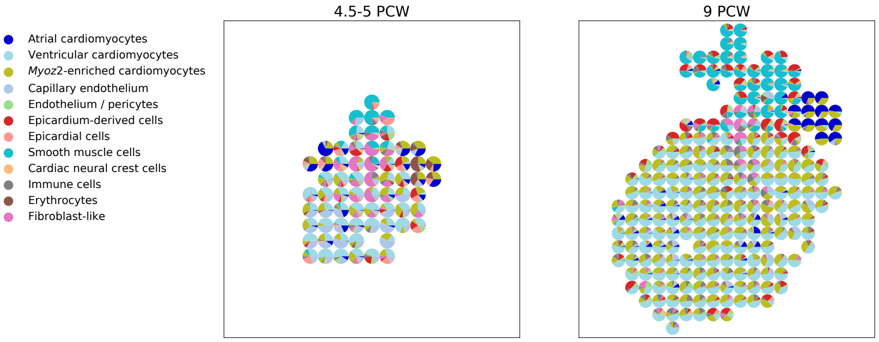


Supplementary Fig. 14: **Pie charts of STitch3D's cell-type deconvolution results for the 4.5-5 PCW and 9 PCW human hearts.** Estimated proportions of cell types obtained from STitch3D in a 4.5-5 PCW heart slice (left) and a 9 PCW heart slice (right) visualized by pie charts.

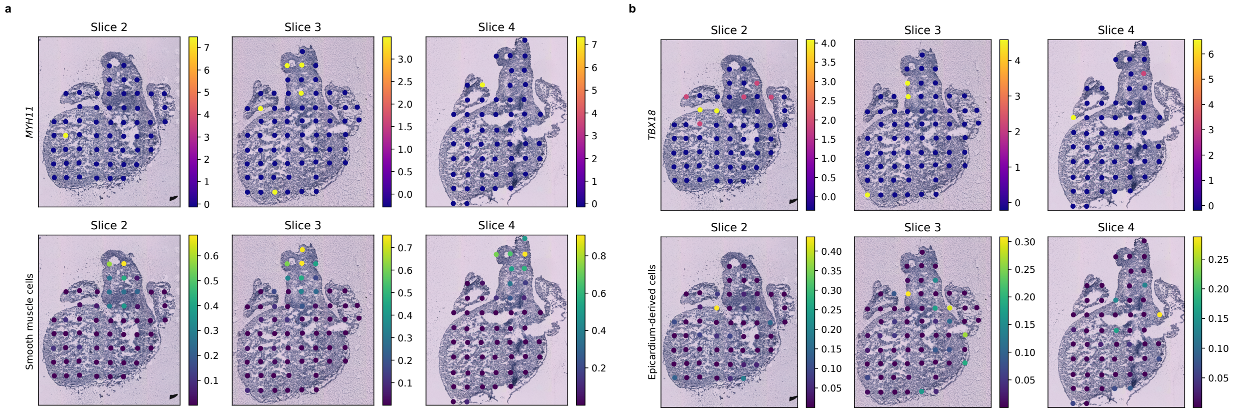


Supplementary Fig. 15: **Comparison between marker genes and STitch3D's cell-type deconvolution result for the 4.5-5 PCW human heart. a**. Expression pattern of smooth muscle-related marker gene *MYH11* (the first row) and STitch3D's estimated cell-type proportions of smooth muscle cells (the second row) visualized on three ST slices. **b**. Expression pattern of EPDC marker gene *TBX18* (the first row) and STitch3D's estimated cell-type proportions of EPDCs (the second row) visualized on three ST slices.

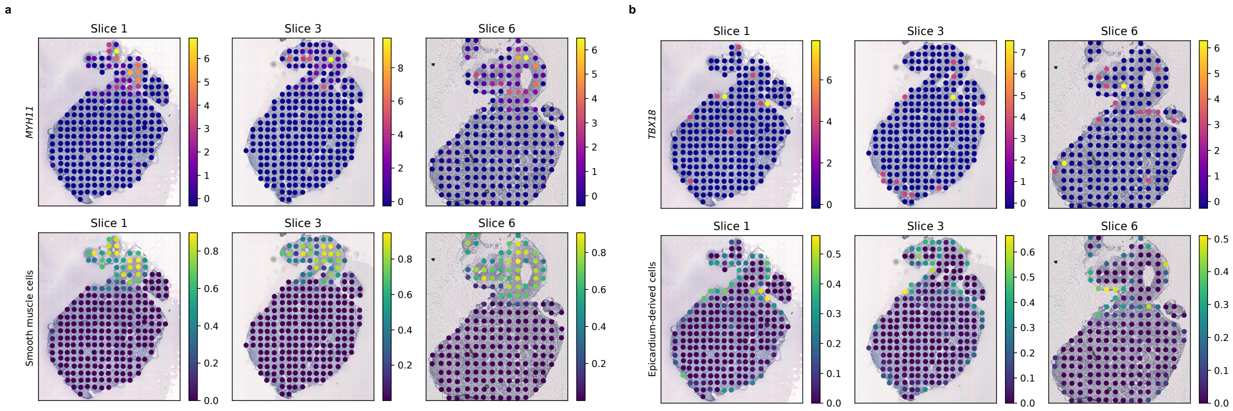


Supplementary Fig. 16: **Comparison between marker genes and STitch3D's cell-type deconvolution result for the 9 PCW human heart. a**. Expression pattern of smooth muscle-related marker gene *MYH11* (the first row) and STitch3D's estimated cell-type proportions of smooth muscle cells (the second row) visualized on three ST slices. **b**. Expression pattern of EPDC marker gene *TBX18* (the first row) and STitch3D's estimated cell-type proportions of EPDCs (the second row) visualized on three ST slices.

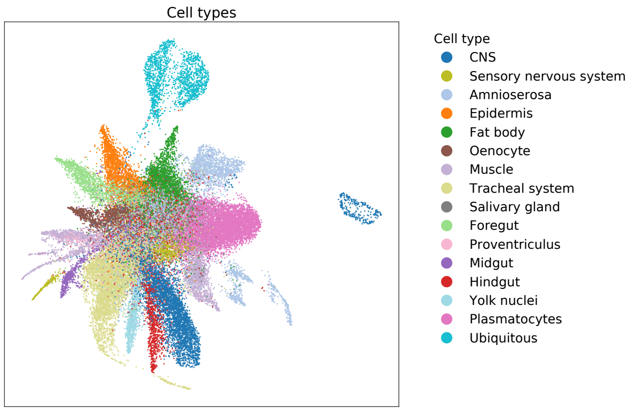


Supplementary Fig. 17: **The UMAP plot of *Drosophila* embryo cells colored by identified regional cell types.** We generated regional cell-type annotation for the sci-RNA-seq *Drosophila* embryo dataset [20].

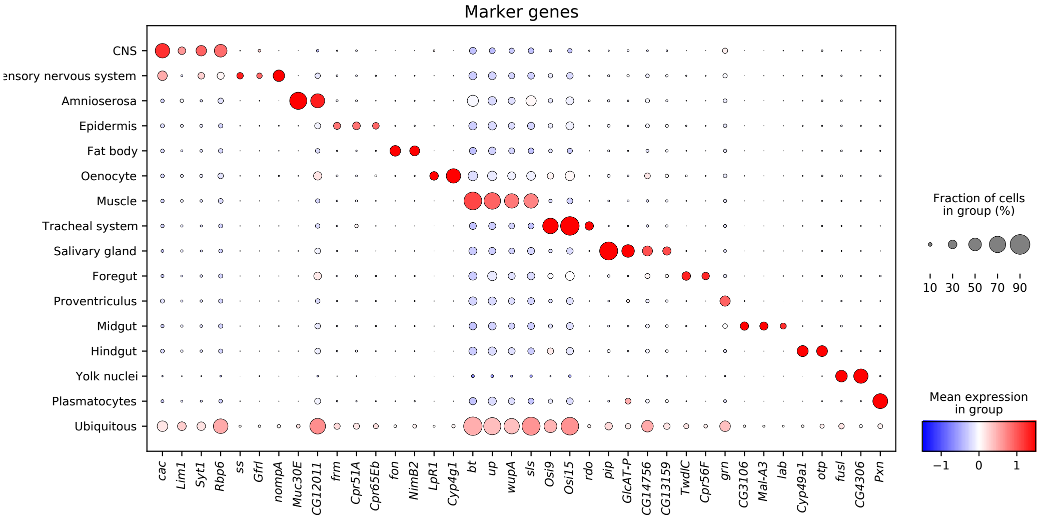


Supplementary Fig. 18: **Marker gene patterns in the sci-RNA-seq *Drosophila* embryo dataset.** The marker genes of regional cell types provided by the Berkeley Drosophila Genome Projec (BDGP) [21, 22] are summarized in Supplementary Table 1.

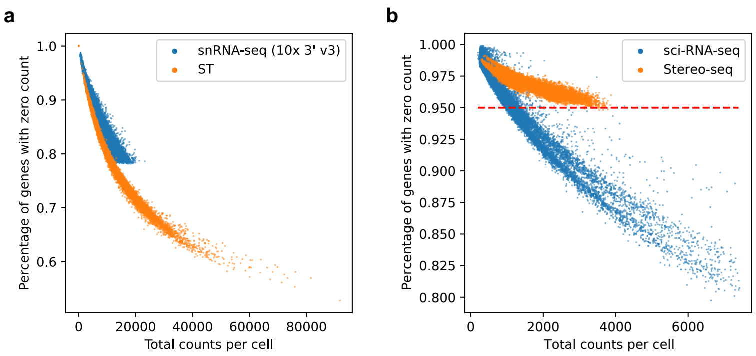


Supplementary Fig. 19: **Sparsity of measured gene expressions in scRNA-seq and ST datasets.** **a**. We visualized the sparsity of mouse brain datasets profiled by snRNA-seq (10x 3' v3) and ST respectively. **b**. We visualized the sparsity of *Drosophila* embryo datasets profiled by sci-RNA-seq and Stereo-seq respectively. For each cell, the $x$ axis shows the total count of genes, and the $y$ axis shows the percentages of genes with zero counts.

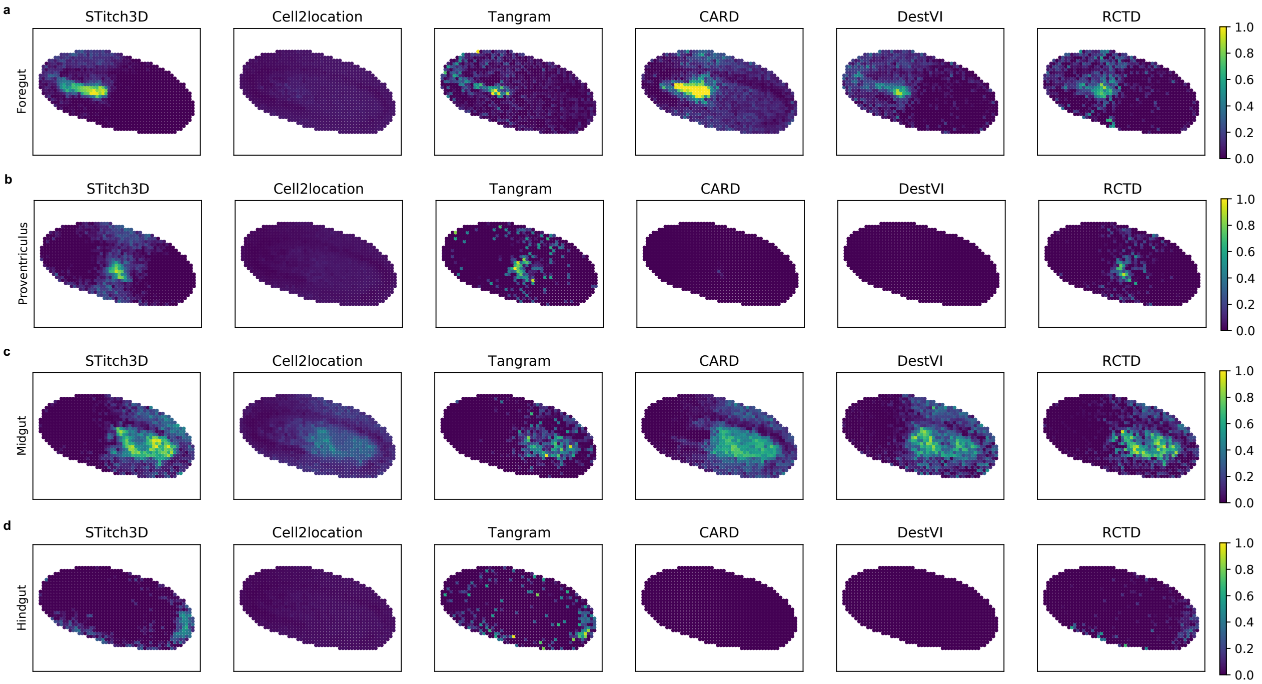


Supplementary Fig. 20: **Comparison of regional cell-type deconvolution results among compared methods.** We visualized estimated proportions of foregut (**a**), proventriculus (**b**), midgut (**c**) and hindgut (**d**) in 2D slices of the *Drosophila* embryo.

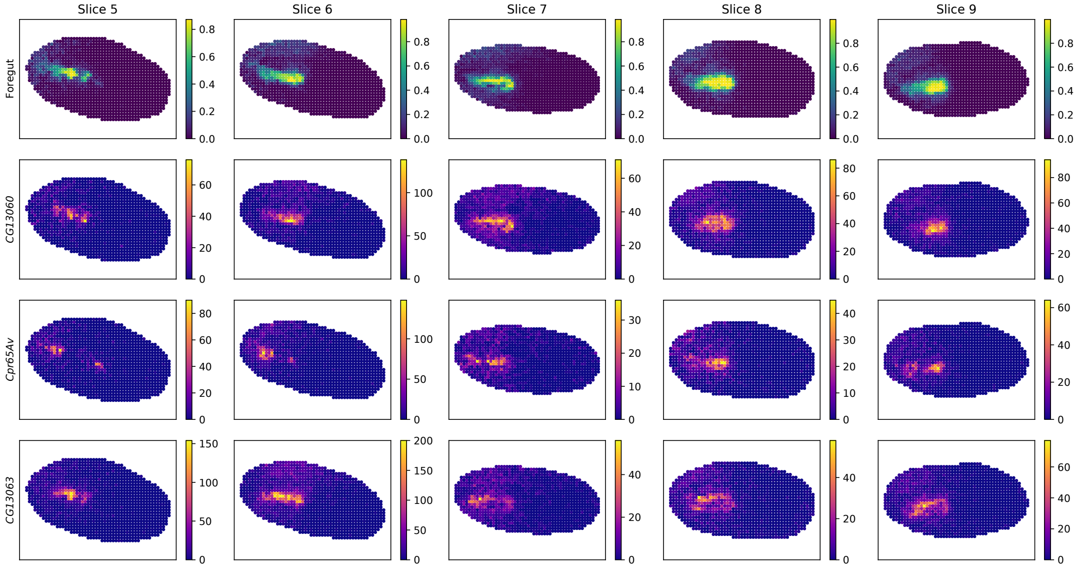


Supplementary Fig. 21: **STitch3D's estimated proportion of foregut, and expression pattern of STitch3D's identified markers of foregut.** We found that *CG13060* was highly expressed in the forgut, *Cpr65Av* and *CG13063* were highly expressed in subregions of the forgut.

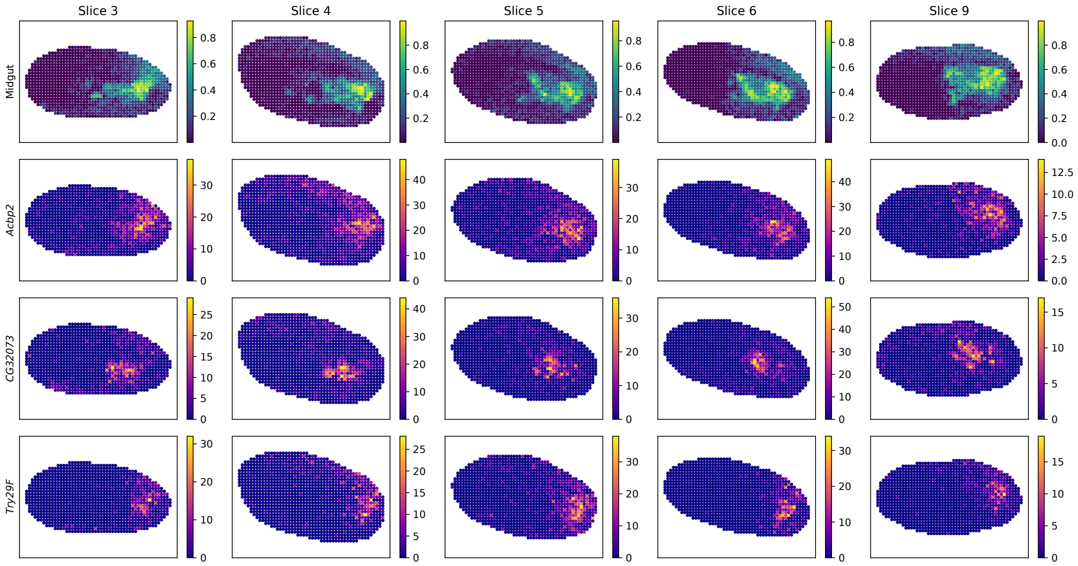


Supplementary Fig. 22: **STitch3D's estimated proportion of midgut, and expression pattern of STitch3D's identified markers of midgut.** We found that *Acbp2* was highly expressed in the midgut, *CG32073* and *Try29F* were highly expressed in subregions of the midgut.

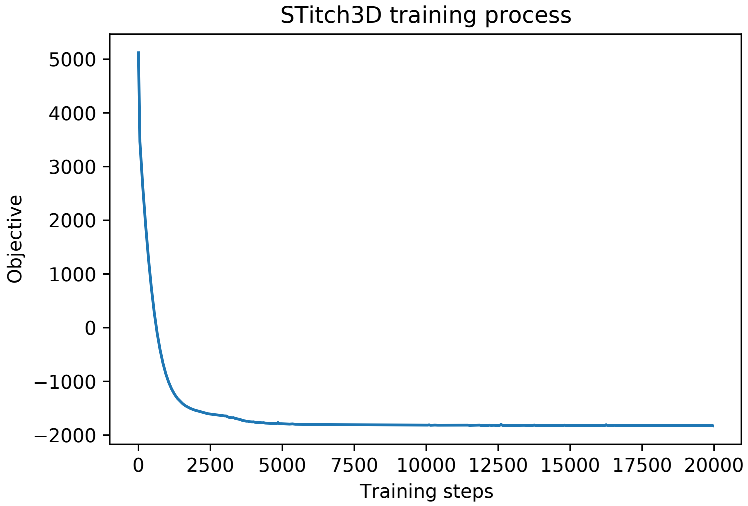


Supplementary Fig. 23: **Convergence of STitch3D's model training.** We recorded STitch3D's training loss versus training steps when applied to the 3D reconstruction of mouse brain.

### Supplementary Tables

| Regional cell type | Genes |
|---|---|
| CNS | *cac*, *Lim1*, *Syt1*, *Rbp6* |
| Sensory nervous system | *ss*, *Gfrl*, *nompA* |
| Amnioserosa | *Muc30E*, *CG12011* |
| Epidermis | *frm*, *Cpr51A*, *Cpr65Eb* |
| Fat body | *fon*, *NimB2* |
| Oenocyte | *LpR1*, *Cyp4g1* |
| Muscle | *bt*, *up*, *wupA*, *sls* |
| Tracheal system | *Osi9*, *Osi15*, *rdo* |
| Salivary gland | *pip*, *GlcAT-P*, *CG14756*, *CG13159* |
| Foregut | *TwdlC*, *Cpr56F* |
| Proventriculus | *grn* |
| Midgut | *CG3106*, *Mal-A3*, *lab* |
| Hindgut | *Cyp49a1*, *otp* |
| Yolk nuclei | *fusl*, *CG4306* |
| Plasmatocytes | *Pxn* |

Supplementary Table 1: **Marker genes for annotating regional cell types in the *Drosophila* embryo [21, 22, 23].**

| ST dataset | # slices | # spots | # cell types | # genes | Time (min) |
|---|---|---|---|---|---|
| Human DLPFC [14] | 4 | 14,243 | 18 | 4,558 | 62.11 |
| Mouse brain [24] | 35 | 17,086 | 59 | 6,227 | 90.85 |
| 4.5-5 PCW human heart [15] | 4 | 232 | 12 | 3,837 | 5.22 |
| 6.5 PCW human heart [15] | 9 | 1,470 | 12 | 3,837 | 7.03 |
| 9 PCW human heart [15] | 6 | 1,340 | 12 | 3,837 | 6.34 |
| *Drosophila* embryo [25] | 13 | 14,132 | 16 | 5,841 | 78.64 |

Supplementary Table 2: **Number of slices, number of spots, number of cell types, number of highly variable genes and STitch3D's corresponding training time for different experiments.**
